# NDUFAB1 Protects Heart by Coordinating Mitochondrial Respiratory Complex and Supercomplex Assembly

**DOI:** 10.1101/302281

**Authors:** Tingting Hou, Rufeng Zhang, Chongshu Jian, Wanqiu Ding, Yanru Wang, Qi Ma, Xinli Hu, Heping Cheng, Xianhua Wang

**Affiliations:** State Key Laboratory of Membrane Biology, Beijing Key Laboratory of Cardiometabolic Molecular Medicine, Peking-Tsinghua Center for Life Sciences, Institute of Molecular Medicine, Peking University, China

**Keywords:** Cardiac protection/Mitochondrial Bioenergetics/NDUFAB1/Respiratory supercomplex

## Abstract

The impairment of mitochondrial bioenergetics, often coupled with exaggerated reactive oxygen species (ROS) production, is emerging as a common mechanism in diseases of organs with a high demand for energy, such as the heart. Building a more robust cellular powerhouse holds promise for protecting these organs in stressful conditions. Here, we demonstrate that NDUFAB1 (NADH:ubiquinone oxidoreductase subunit AB1), acts as a powerful cardio-protector by enhancing mitochondrial energy biogenesis. In particular, NDUFAB1 coordinates the assembly of respiratory complexes I, II, and III and supercomplexes, conferring greater capacity and efficiency of mitochondrial energy metabolism. Cardiac-specific deletion of *Ndufab1* in mice caused progressive dilated cardiomyopathy associated with defective bioenergetics and elevated ROS levels, leading to heart failure and sudden death. In contrast, transgenic overexpression of *Ndufab1* effectively enhanced mitochondrial bioenergetics and protected the heart against ischemia-reperfusion injury. Our findings identify NDUFAB1 as a central endogenous regulator of mitochondrial energy and ROS metabolism and thus provide a potential therapeutic target for the treatment of heart failure and other mitochondrial bioenergetics-centered diseases.

## Introduction

Being so-called “powerhouses”, mitochondria comprise up to 40% of the heart mass and are central to cardiac bioenergetics (Ingwall, 2002, Neubauer, 2007). As hazardous by-products of energy metabolism, reactive oxygen species (ROS) are also emitted from the respiratory electron-transfer chain (ETC), and excessive ROS emission imposes oxidative stress to damage proteins, lipids, and even genetic materials (Beckman & Ames, 1998, Droge, 2002). Studies have shown that >50% of individuals with mutations in genes encoding mitochondrial proteins eventually develop cardiomyopathy (DiMauro & Schon, 2003), suggesting that defective cardiac bioenergetics as well as its interlinked ROS metabolism is a fundamental disease mechanism in the heart (Ingwall, 2004, Maack & O’Rourke, 2007). As such, building a more robust cellular powerhouse with decreased ROS emission holds promise for protecting the heart in stressful conditions.

The most important player in mitochondrial energy production is the ETC, which consists of four multi-heteromeric complexes (complexes I–IV) in the inner membrane of the organelle. They catalyze outward proton movement to build a transmembrane electrochemical gradient that drives ATP synthase (complex V) for ATP synthesis (Nicholls & Ferguson, 2002). Individual ETC complexes can organize into supercomplexes (SCs) of different composition and stoichiometry, which have recently been visualized by cryo-electron microscopy (Gu et al., 2016, Guo et al., 2017, Letts et al., 2016, Sousa et al., 2016, Wu et al., 2016). Such SCs are thought to channel electron transfer more efficiently, limit ROS production, protect vulnerable ETC sites in the complexes from oxidative damage, and stabilize individual complexes (Acin-Perez et al., 2004, Diaz et al., 2006, Letts et al., 2016, Lopez-Fabuel et al., 2016, Maranzana et al., 2013, Milenkovic et al., 2017, Porras & Bai, 2015, Schagger et al., 2004). Interestingly, SC formation is dynamically regulated in accordance with changes in cellular metabolism (Acin-Perez & Enriquez, 2014, Greggio et al., 2017, Guaras et al., 2016, Lapuente-Brun et al., 2013), and deficiency of SCs is associated with heart failure (Rosca et al., 2008). Therefore, enhancing SC formation might afford an effective therapeutic strategy to make the mitochondria more robust and efficient powerhouses.

NDUFAB1 (NADH:ubiquinone oxidoreductase subunit AB1), also known as mitochondrial acyl carrier protein (Runswick et al., 1991), not only participates in the synthesis of lipoic acid in the type II fatty acid biosynthetic (FASII) pathway (Feng et al., 2009, Hiltunen et al., 2010b), but also acts as an accessory subunit of complex I with a 2:1 stoichiometry (Vinothkumar et al., 2014). A recent study has shown that NDUFAB1 interacts with seven mitochondrial LYR motif-containing (LYRM) proteins: LYRM1, LYRM2, LYRM4/ISD11, LYRM5, LYRM7/MZM1L, SDHAF3/ACN9, and FMC1/C7orf55 (Floyd et al., 2016). Most are linked to various mitochondrial metabolic processes, such as the assembly of complex II (for SDHAF3/ACN9) and complex V (for FMC1/C7orf55), deflavination of the electron-transferring flavoprotein (for LYRM5), and biosynthesis of iron-sulfur (FeS) centers (for LYRM4/ISD11) (Floyd et al., 2016). In yeast cells, Van *et al* have shown that NDUFAB1 interacts with and stabilizes the FeS biogenesis complex (Van Vranken et al., 2016). Thus, NDUFAB1 might be a core player in orchestrating mitochondrial energy biogenesis.

In this study, we aimed to test the potential protective role of NDUFAB1 in the mammalian heart using cardiac-specific *Ndufab1* knockout (cKO) and transgenic overexpression (TG) mouse models. We found that NDUFAB1 coordinates the assembly of individual ETC complexes and SCs, acting as a central endogenous regulator of the efficiency and capacity of mitochondrial energy metabolism as well as ROS emission. More importantly, NDUFAB1 exerts a cardio-protective effect when the heart is subjected to ischemia-reperfusion (IR) injury. Our findings substantiate that targeting the mitochondria for greater bioenergetic capacity and repressed ROS emission is an effective strategy for cardio-protection, and mark NDUFAB1 as a potential therapeutic target for the treatment of cardiomyopathy.

## Results

### Cardiac-specific ablation of NDUFAB1 causes progressive dilated cardiomyopathy leading to heart failure

NDUFAB1 was widely expressed in different tissues and was particularly enriched in the heart (Fig EV1A). Whole-body knockout of *Ndufab1* was embryonic lethal (Fig EV1B and 1C), showing that NDUFAB1 is essential for developmental viability. To explore the potential cardiac functions of NDUFAB1, we generated a cardiac-specific knockout mouse model (cKO) wherein exon 3 of *Ndufab1* was flanked by loxP sites (*Ndufab1*^flox/flox^, Fig EV1B). The *Ndufab1*^flox/flox^ mice were cross-bred with Mlc2v-Cre mice to allow cardiomyocyte-specific deletion of *Ndufab1*, as previously described (Zheng et al., 2009). As shown in Figure 1A and Figure EV1D, cardiac expression of *Ndufab1* was decreased by ~90% at the protein level in cKO mice compared to their wild-type (WT) littermates. The cKO mice were smaller, had a reduced body weight, and even started to lose weight precipitously after 14 weeks of age (Fig 1B). Meanwhile, the lifespan of cKO mice was markedly shortened, sudden death beginning at ~12 weeks of age; the maximal lifespan was <19 weeks (Fig 1C).

**Figure 1.**
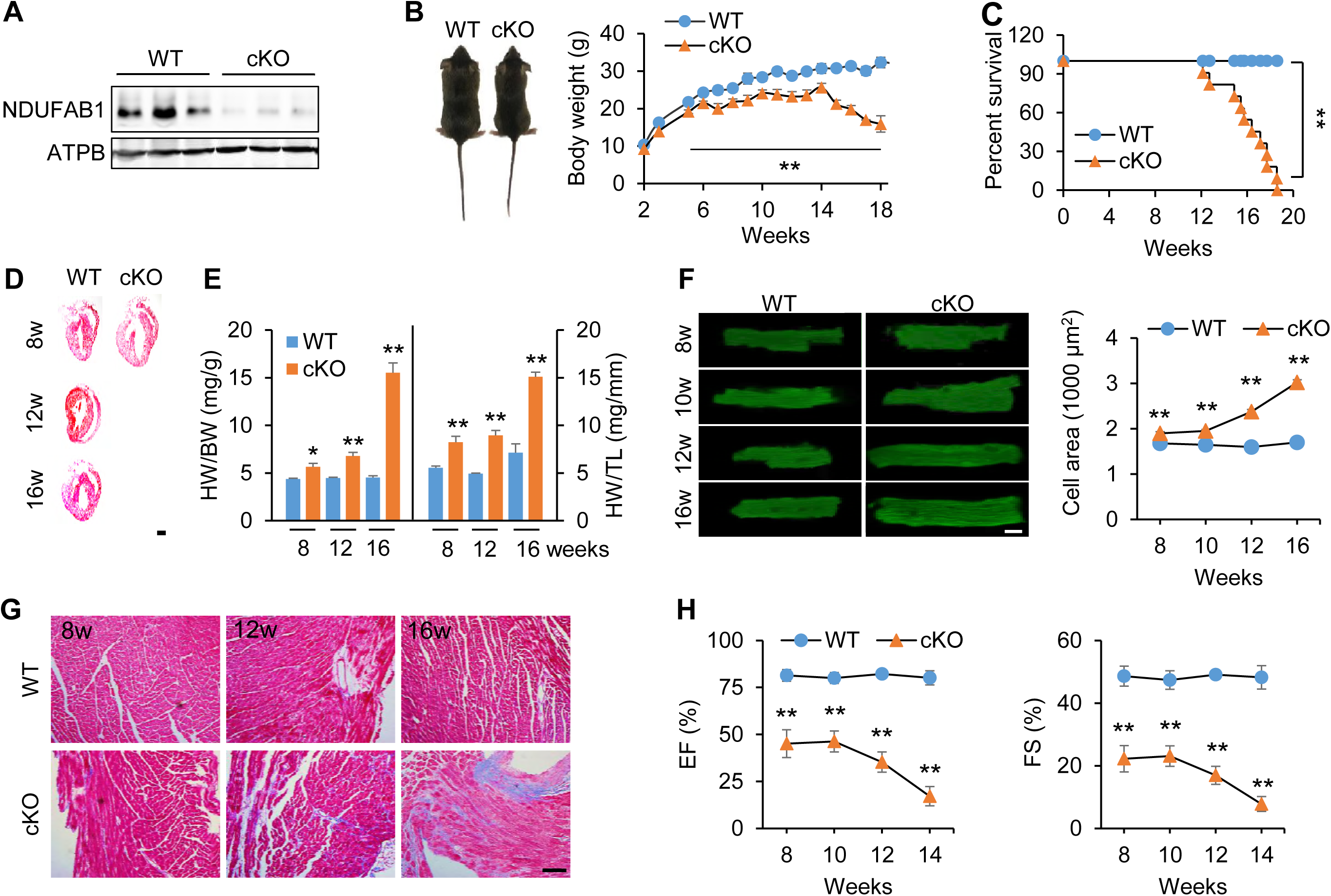
Cardiac-specific knockout of *Ndufab1* (cKO) causes dilated cardiomyopathy leading to heart failure in mice. **(A)** Western-blots of NDUFAB1 in wild-type (WT) and cKO mouse hearts. Anti-ATPB served as the loading control. **(B)** Effect of cardiac NDUFAB1 ablation on mouse body weight (mean ± s.e.m.; n = 3–16 male mice per group per time point; ^∗∗^***P*** <0.01 *versus* WT at the same time point). Inset, photograph of WT and cKO mice at 14 weeks of age. **(C)** Kaplan-Meier survival curves for WT and cKO mice (mean ± s.e.m.; n = 11 male mice per group; ^∗∗^ ***P*** <0.01 *versus* WT). **(D)** Representative longitudinal sections of hearts from male mice 8–16 weeks (w) old. Scale bar, 5 mm. **(E)** Ratios of heart weight (HW) to body weight (BW) or tibia length (TL) (mean ± s.e.m.; n = 3–14 male mice per group; ^∗^***P*** <0.05, ^∗∗^***P*** <0.01 *versus* WT at the same age). **(F)** Progressive hypertrophy of cardiomyocytes from cKO mice. Left: representative confocal micrographs of cardiomyocytes stained with DCF (scale bar, 20 μm). Right: statistics (mean ± s.e.m.; n = 281–611 cells from 4 male mice per group; ^∗^***P*** <0.05, ^∗∗^***P*** <0.01 *versus* WT at the same age). (G) Fibrosis of left ventricular tissue (Masson’s trichrome staining; scale bar, 100 μm). (H) Echocardiographic analysis of cardiac function; EF, ejection fraction; FS, fractional shortening (mean ± s.e.m.; n = 3–9 male mice per group; ^∗∗^***P*** <0.01 *versus* WT at the same age).

We next examined the effect of NDUFAB1 ablation on cardiac morphology and functions at different ages. The cKO hearts progressively enlarged with age and developed dilated cardiomyopathy (Fig 1D and Fig EV1E). Compared to littermates, the heart weight/body weight ratio in cKO mice was increased by 26.9% at 8 weeks, 51.5% at 12 weeks, and 241.7% at 16 weeks; and the heart weight/tibia length ratio was increased by 45.5% at 8 weeks, 81.0% at 12 weeks, and 111.3% at 16 weeks (Fig 1E). At the cellular level, the longitudinal-section area of cardiomyocytes was nearly constant in WT but increased progressively in cKO mice, reaching a 78% increase at 16 weeks (Fig 1F), indicative of cardiomyocyte hypertrophy. Masson staining showed that NDUFAB1 ablation induced extensive cardiac fibrosis early, at 8 weeks (Fig 1G and Fig EV1F). Echocardiography revealed that the EF and FS of cKO hearts were halved at this age, and EF diminished by 79% and FS by 84% at the age of 14 weeks (Fig 1H), while the heart rate remained unchanged (Fig EV2A). The thickness of both the left ventricular posterior wall and the inter-ventricular septum decreased while the left ventricular diameter and volume increased with age in cKO hearts (Fig EV2B and 2C). These findings indicate that cardiac-specific ablation of NDUFAB1 leads to dilated cardiomyopathy accompanied by cardiomyocyte hypertrophy, interstitial fibrosis, and systolic and diastolic dysfunction, eventually resulting in heart failure and sudden death, revealing an essential role of NDUFAB1 in cardiac function.

### Mitochondrial dysfunctions in *Ndufab1* cKO heart

To investigate the mechanism underlying the cardiomyopathy and heart failure induced by NDUFAB1 ablation, we next assessed changes in cardiac mitochondrial functions in this genetic model of cardiac disease. Electron microscopic analysis revealed that, in 16 week-old cKO mice, cardiac mitochondria were in disarray along the sarcomeres with some exhibiting structural damage and deformities (Fig 2A), while the individual mitochondrial cross-sectional area was unaltered and the total mitochondrial volume fraction was slightly increased (Fig EV3). Measuring mitochondrial membrane potential (ΔΨ_m_) in isolated cardiomyocytes with the potential-sensitive fluorescent probe tetramethyl rhodamine methyl ester (TMRM) showed that NDUFAB1 ablation decreased ΔΨ_m_ at all ages (Fig 2B). Measurements with the fluorescent probe mitoSOX showed that mitochondrial ROS level was significantly elevated in cKO cardiomyocytes from 10-week-old mice compared to WT, and reached a 88% increase of mitoSOX fluorescence at 14-16 weeks (Fig 2D). However, the cellular ATP level, as measured with the luciferin luminescence assay, remained unchanged in 8-week-old mice despite the impairment of mitochondrial function, but was significantly lowered in 14-16-week or older cKO mice (Fig 2C). This result suggests that, while the ATP level is initially tightly safeguarded in the heart (Allue et al., 1996, Balaban et al., 1986, Matthews et al., 1981, Neely et al., 1973, Wang et al., 2017), NDUFAB1 ablation curtails the energy reserve capacity, exhaustion of which impairs ATP homeostasis. Taken together, these results indicate that NDUFAB1 acts as a central endogenous regulator for mitochondrial bioenergetics and ROS metabolism.

**Figure 2.**
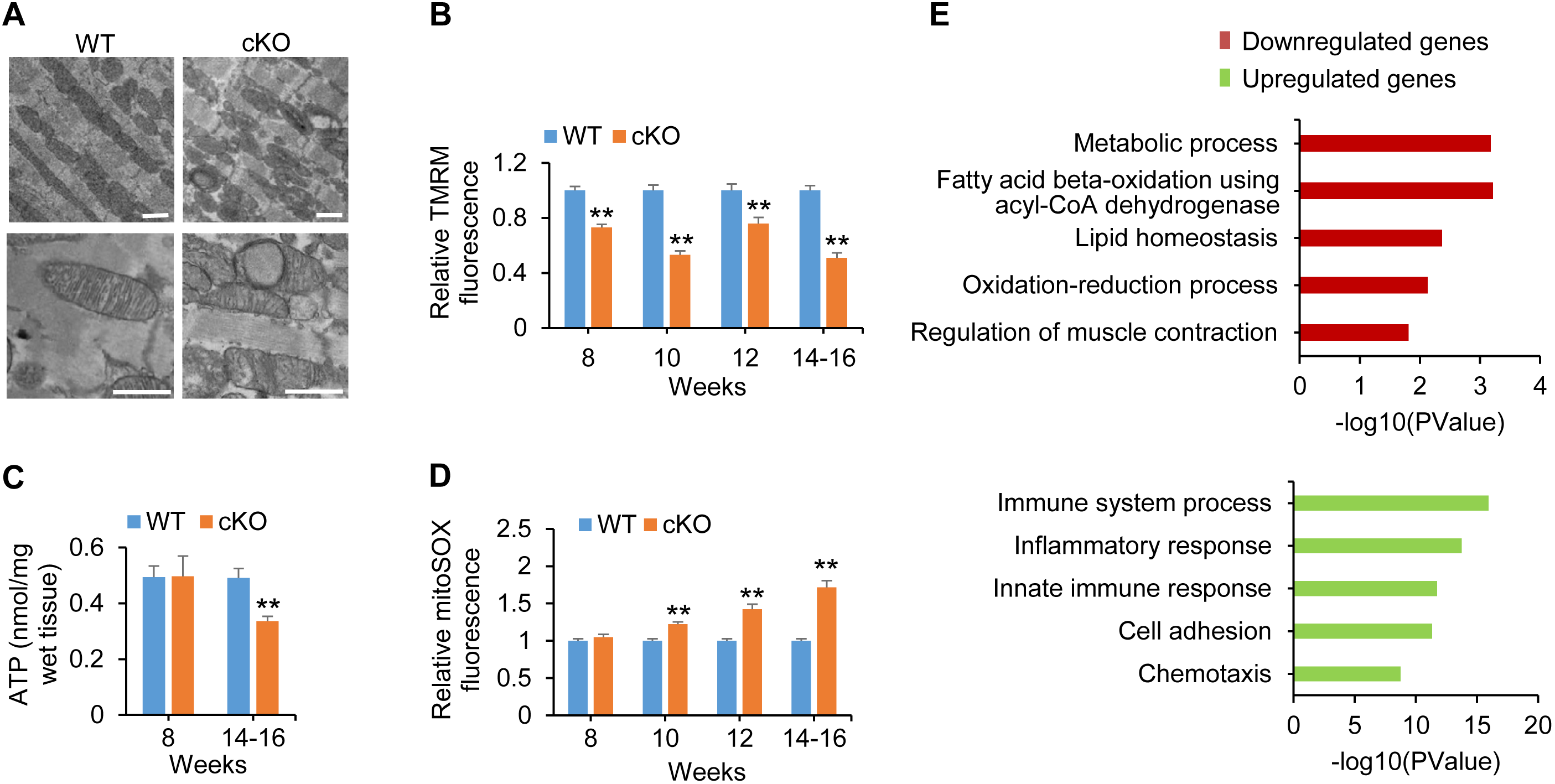
Defective mitochondrial bioenergetics and ROS metabolism in *Ndufab1* cKO mice. **(A)** Electron micrographs of mitochondria in left ventricular tissue from 16-week old male mice (scale bars, 1 μm). **(B)** Decline of ΔΨ_m_ in cKO myocardium assessed by TMRM staining and confocal imaging of isolated cardiomyocytes. The cKO TMRM fluorescence was normalized to WT at the same age (n = 20–58 cells from 4 male mice per group; *∗∗**P*** <0.01 *versus* WT). **(C)** Cardiac ATP levels measured with luciferin assay (n = 5–9 male mice per group; ^∗∗^***P*** <0.01 *versus* WT). **(D)** Progressive elevation of mitochondrial ROS measured with mitoSOX. The cKO mitoSOX fluorescence was normalized to WT at the same age (n = 102–309 cells from 4–5 male mice per group; ^∗∗^***P*** <0.01 *versus* WT). **(E)** Top 5 enriched Gene Ontology terms of downregulated and upregulated genes. RNA-seq analysis was performed on cKO and WT hearts (n = 4 male mice per group) and the genes upregulated or downregulated were analyzed.

To delineate molecular alterations associated with NDUFAB1 ablation, we analyzed the transcriptome of left ventricles from cKO and WT mice of 8-16 weeks by RNA sequencing (RNA-seq). Among 11494 genes analyzed, 430 were downregulated and 1081 were upregulated. Interestingly, the top 5 Gene Ontology terms of the downregulated genes were mainly associated with mitochondrial metabolism, including metabolic process, fatty acid metabolism, lipid homeostasis, oxidation-reduction process (Fig 2E, Tables EV1 and EV2), in good agreement with the defective bioenergetics phenotype of the cKO heart (Fig 2B-2D). The downregulated genes involving regulation of muscle contraction is consistent with the phenotype of cardiomyopathy (Fig 2E). Meanwhile, the top upregulated Gene Ontology terms were mostly involved in immune system process, inflammatory response, cell adhesion, and chemotaxis (Fig 2E, Tables EV3 and EV4). We deduce that these upregulated processes might reflect compensatory as well as maladaptive responses to cardiac damage due to mitochondrial dysfunctions. These changes in the transcriptome underscore that NDUFAB1 is a nodal point in coordinating various processes of mitochondrial energy metabolism.

### Impaired assembly of mitochondrial ETC complexes and SCs in cKO hearts

The multifaceted dysfunctions of mitochondrial energy and ROS metabolism in NDUFAB1 cKO hearts suggest a promiscuous mechanism of action for this mitochondrial protein. In this regard, NDUFAB1 is found not only as a subunit of complex I but also to participate in synthesis of lipoic acid in FASII pathway (Feng et al., 2009, Hiltunen et al., 2010b) and in FeS biogenesis in yeast (Cory et al., 2017, Van Vranken et al., 2016). We first tested the changes of lipoic acid by measuring pyruvate dehydrogenase activity as lipoic acid is its obligate cofactor (Hiltunen et al., 2010a). The similar pyruvate dehydrogenase activity between WT and cKO hearts indicates NDUFAB1 ablation did not affect FASII pathway in the heart (Fig EV4). We therefore investigated whether and how NDUFAB1 ablation impacts on the assembly and activity of individual ETC complexes and supercomplexes in the mammalian heart. First, we measured the respiratory activity of isolated mitochondria in the presence of different substrates. In the presence of complex I substrate, the oxygen consumption rate (OCR) of cKO mitochondria was significantly decreased (Fig 3A), revealing an important functional role of NDUFAB1 as an accessory subunit of complex I (Vinothkumar et al., 2014). Interestingly, the cKO mitochondria also displayed a decreased OCR when complex II or complex III substrates were added (Fig 3A), whereas the OCR supported by complex IV substrate was not significantly altered (Fig 3A). These data suggest that, rather than merely a complex I subunit, NDUFAB1 alters the function of the ETC at multiple sites.

**Figure 3.**
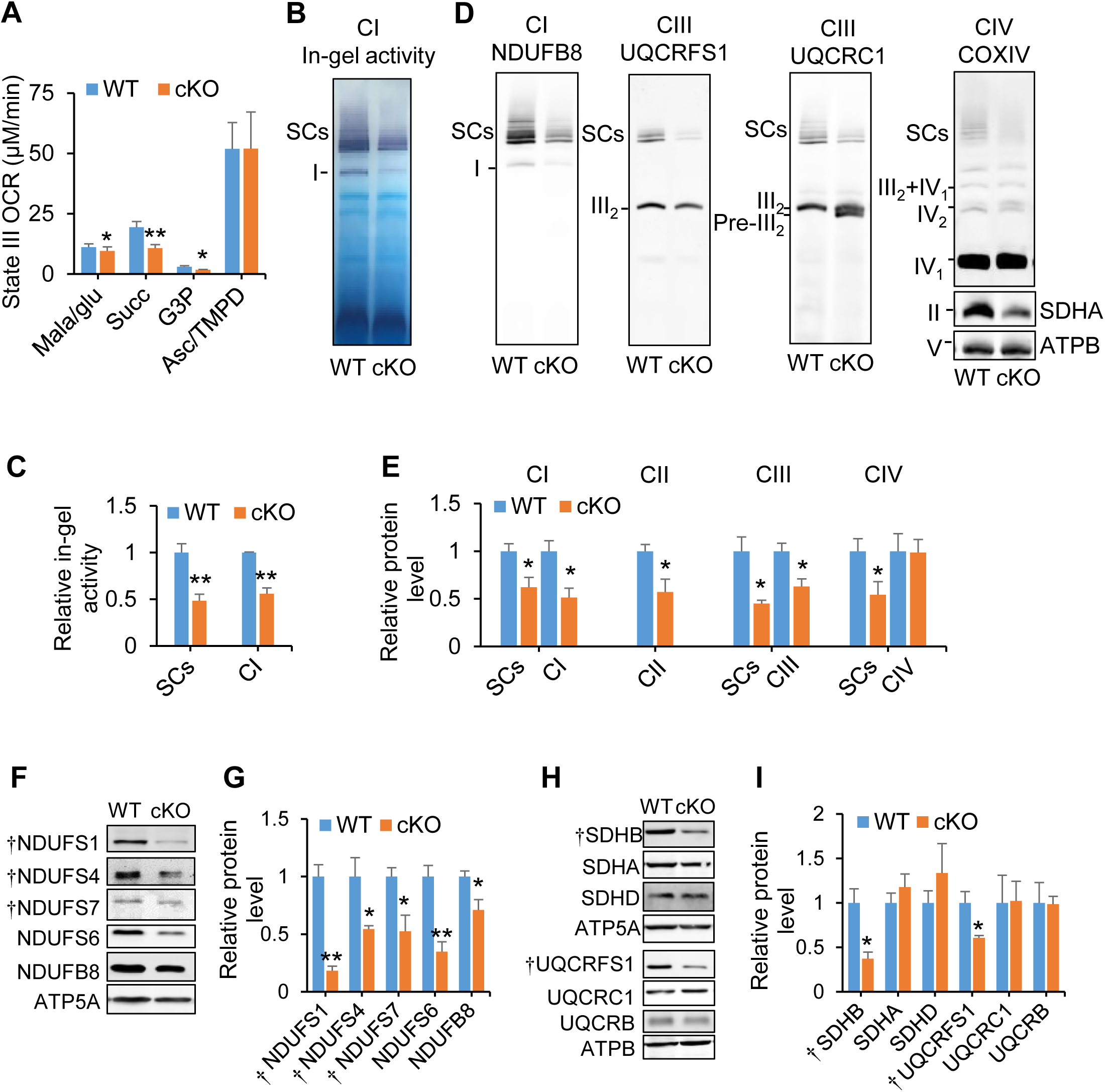
Defective respiration and impaired assembly of ETC complexes I-III and SCs in cKO hearts. **(A)** Effect of NDUFAB1 ablation on state III oxygen consumption rate (OCR) in isolated mitochondria. Different respiratory substrates were used: malate/glutamate (Mala/glu, 5 mM) for complex I, succinate (Succ, 5 mM) for complex II, glycerol-3-phosphate (G3P, 5 mM) for complex III, and ascorbate (Asc, 2.5 mM)/N,N,N’,N’-tetramethyl-p-phenylenediamine (TMPD, 0.5 mM) for complex IV, and ADP (100 μM) (mean ± s.e.m.; n = 3–14 male mice of 10-week old per group; ^∗^***P*** <0.05, *∗∗**P***<0.01 *versus* WT). **(B)** In-gel activity of SCs and complex I. **(C)** Statistics of **(B)**. The cKO activity was normalized to WT (mean ± s.e.m.; *n* = 3–4 male mice per group; ^∗∗^***P*** <0.01 *versus* WT). **(D)** BN-PAGE immunoblots of individual ETC complexes and SCs. SCs were visualized by antibodies against subunits of complex I (CI, NDUFB8), complex III (CIII, UQCRFS1 or UQCRC1), and complex IV (CIV, COX IV). Complex II was visualized by antibody against SDHA, and complex V with antibody against ATPB. **(E)** Statistics of **(D)**. The complex and supercomplex expression of cKO was normalized to WT (mean ± s.e.m.; *n* = 3–5 independent experiments per group; ^∗^***P*** <0.05 *versus* WT). Anti-ATPB served as the loading control; for CIII, anti-UQCRFS1 blots were used. **(F)** Western blots of complex I subunits. ^†^FeS-containing subunits; anti-ATP5A served as the loading control. **(G)** Statistics of **(F)**. The protein level of cKO was normalized to WT (mean ± s.e.m.; *n* = 3–5 independent experiments per group; ^∗^***P*** <0.05, ^∗∗^***P*** <0.01 *versus* WT). **(H)** Western blots of subunits of complexes II and III. ^†^FeS-containing subunits. **(I)** Statistics of **(H)**. The protein level of cKO was normalized to WT (mean ± s.e.m.; *n* = 3–5 independent experiments per group; ^∗^***P*** <0.05 *versus* WT). Male mice used in **(B)-(I)** were 16 weeks old.

Further, we assessed the assembly of ETC complexes using blue native polyacrylamide gel electrophoresis (BN-PAGE). Consistent with previous reports (Jian et al., 2017, Mitsopoulos et al., 2015), complexes I, III, and IV were assembled into SCs, as evidenced by immunoblot analysis following BN-PAGE using antibodies against NDUFB8 (complex I), SDHA (complex II), UQCRFS1 or UQCRC1 (complex III), COXIV (complex IV), and ATP5A (complex V), while complexes II and V mainly remained as individual entities in both WT and cKO groups (Fig 3D). Strikingly, NDUFAB1 ablation decreased the SCs containing complexes I, III, and IV by 38%, 55%, and 46% respectively (Fig 3D and 3E), indicating a crucial role of NDUFAB1 in SC formation. We also assessed the abundance of individual complexes and found that individual complexes I, II, and III were remarkably diminished (Fig 3D and 3E) while complex IV was not significantly changed in cKO mitochondria (Fig 3D and 3E), consistent with the aforementioned functional data (Fig 3A). Meanwhile, in-gel activity analysis in the presence of complex I substrate revealed that the activity of either complex I or SCs was markedly lowered in the absence of NDUFAB1 (Fig 3B and 3C). Thus, we conclude that NDUFAB1 coordinates the assembly and activity of individual complexes I, II, and III as well as SCs, thereby regulating mitochondrial energy biogenesis.

We noted that a complex III assembly intermediate lacking UQCRFS1 (ubiquinol-cytochrome c reductase, Rieske iron-sulfur polypeptide 1), which is an FeS-containing subunit and is incorporated into complex III at the last step (Wagener et al., 2011), accumulated in cKO mitochondria (Fig 3D). This finding hinted at the possibility that NDUFAB1 regulates the assembly of ETC complexes and SCs through its ability to regulate FeS biogenesis (Cory et al., 2017, Van Vranken et al., 2016). Indeed, western-blot analysis showed that only the FeS-containing subunits SDHB of complex II and UQCRFS1 of complex III were significantly decreased while the other subunits examined were unaltered in the absence of NDUFAB1 (Fig 3H and 3I); that is, NDUFAB1 ablation selectively disrupts FeS-containing subunits. For complex I, however, we found all subunits examined, whether FeS-containing (NDUFS1, NDUFS4, and NDUFS7) or not (NDUFS6 and NDUFB8), were dramatically decreased in cKO mitochondria (Fig 3F and 3G), suggesting additional regulatory mechanisms exist (see Discussion). Since the mRNA levels of most of the above subunits of complexes I-III were not significantly altered (Fig EV5), NDUFAB1 regulation appears to occur at the protein rather than the mRNA level, presumably by affecting protein stability.

### Enhanced mitochondrial functions in *Ndufab1* transgenic hearts

The results thus far revealed that NDUFAB1 is a key molecule in coordinating the dynamic assembly of individual ETC complexes and SCs and thereby acts as a central regulator of mitochondrial bioenergetics and ROS metabolism. We reckoned that augmenting the abundance of NDUFAB1 might provide an effective strategy to build a more robust and efficient powerhouse for the benefit of the heart, providing that endogenous NDUFAB1 is present at a sub-saturating level. To test this possibility, we generated an *Ndufab1* transgenic mouse model (TG) using a β-actin promoter (Fig EV6A) in which NDUFAB1 was overexpressed ~8-fold in the heart (Fig 4A and Fig EV6B). Compared with WT littermates, TG mice grew normally and had similar lifespans under our experimental conditions (Fig EV6C and 6D). Cardiac morphology and functional performance determined by echocardiography (Fig EV6E-6G) and mitochondrial morphology revealed by electron microscopy (Fig EV6H) were all comparable in WT and TG hearts. Nonetheless, the mitochondrial ROS level was markedly decreased in TG cardiomyocytes (Fig 4C), suggesting an alleviation of basal oxidative stress. The ΔΨ_m_ was significantly increased (Fig 4B), and the maximal OCR measured by Seahorse assay was higher in TG than in WT cardiomyocytes (Fig 4D and 4E). This result implies an enhanced reserve capacity for ATP production, although the homeostatic ATP level was unaltered (Fig EV6I). Moreover, the respiratory control ratio was significantly improved in the presence of substrates of complex I, II, or III (Fig 4F), indicating greater efficiency of mitochondrial energy metabolism.

**Figure 4.**
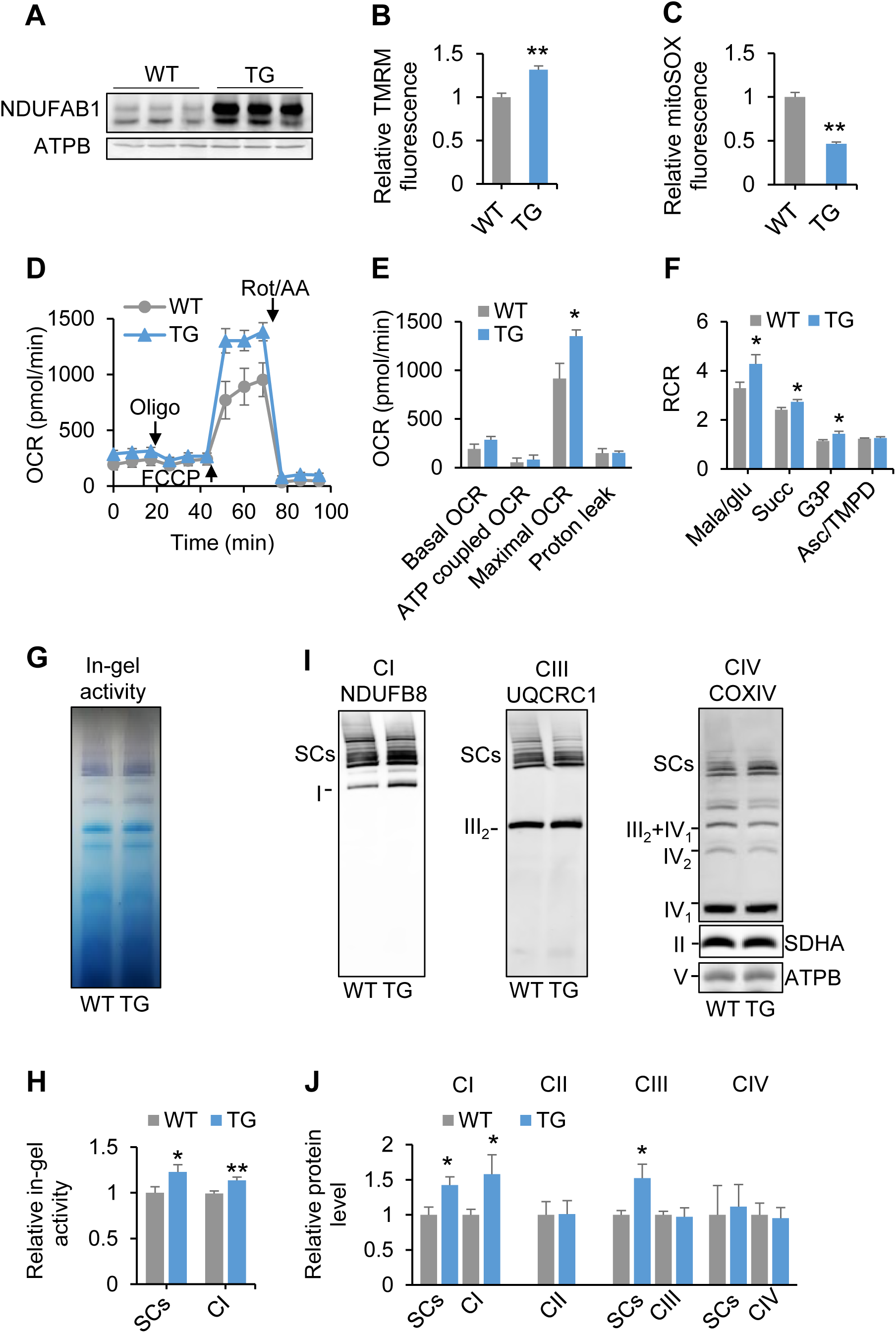
Enhanced mitochondrial bioenergetics and SC assembly in the heart of *Ndufab1* transgenic mice (TG). TG and WT littermates 2–4 months old were used. **(A)** Western blots for NDUFAB1 expression in WT and TG hearts. Anti-ATPB served as the loading control. **(B)** Increased ΔΨ_m_ in cardiomyocytes isolated from TG mice. The TMRM fluorescence of TG was normalized to WT (n = 38–43 cells from 3 male mice per group; ^∗∗^***P*** <0.01 *versus* WT). **(C)** Decreased mitochondrial ROS level in TG cardiomyocytes. The mitoSOX fluorescence of TG was normalized to WT (n = 34–41 cells from 3 male mice; ^∗∗^***P*** <0.01 *versus* WT). **(D)** Whole-cell OCR measured with Seahorse. Arrows indicate the times of adding oligomycin (Oligo, 1 μM), carbonyl cyanide 4-(trifluoromethoxy) phenylhydrazone (FCCP, 0.5 μM), and rotenone/antimycin A (Rot/AA, 1 μM each). **(E)** Statistics of basal, ATP-coupled, maximal, and proton-leak-associated OCR. In **(D)** and **(E)**, data are mean ± s.e.m. *n* = 4–5 male mice per group. ^∗^***P*** <0.05 *versus* WT. **(F)** Changes of respiratory control ratio (RCR) with different substrates in isolated mitochondria (mean ± s.e.m.; n = 3–6 male mice per group; ^∗^***P*** <0.05 *versus* WT). **(G)** In-gel activity of SCs and complex I. **(H)** Statistics of **(G)**. The activity of TG was normalized to WT (mean ± s.e.m.; n = 5 male mice per group; ^∗∗^***P*** <0.01 *versus* WT). **(I)** BN-PAGE immunoblots of SCs and individual ETC complexes as in Figure 3D. **(J)** Statistics of **(I)**. The complex and supercomplex expression of TG was normalized to WT (mean ± s.e.m.; n = 4–8 male mice per group).

At the molecular level, in-gel activity analysis showed that both complex I and SC activity were significantly enhanced in TG hearts (Fig 4G and 4H), and BN-PAGE analysis showed that NDUFAB1 overexpression significantly increased the assembly of complex I and SCs (Fig 4I and 4J), while the expression of all subunits examined, including those for complexes I-III, was unchanged (Fig EV6J and 6K). Thus, NDUFAB1 overexpression promotes the assembly of complex I and SCs, conferring on mitochondria a greater capacity and efficiency of energy metabolism and less ROS emission.

### NDUFAB1 overexpression protects the heart against IR injury

The cardiac performance as well as the lifespan of TG mice was similar to that of WT littermates under unchallenged conditions (Fig EV6). However, TG mice might be more resistant to cardiac injury because of augmented capacity and efficiency of mitochondrial bioenergetics along with attenuated ROS production. Next, we subjected the WT and TG hearts to IR injury to unmask this potential cardio-protective role of NDUFAB1. First, we applied an *ex vivo* IR protocol based on Langendorff perfusion with 30-min ischemia followed by 30-min reperfusion (Fig EV7A). Since excessive ROS production during IR is a major contributor to IR injury (Eltzschig & Eckle, 2011, Murphy & Steenbergen, 2008, Yellon & Hausenloy, 2007), we tracked the ROS production with indicator 5-(and-6)-chloromethyl-2 stadichlorodihydrofluorescein diacetate acetyl ester (DCF) (Fig 5A). In WT hearts, there was a trend of increasing ROS during ischemia followed by a prominent ROS burst immediately after reperfusion (Fig 5A and 5B). Remarkably, the ROS burst after IR was ameliorated by 34% in TG hearts (Fig 5A and 5B). It has recently been reported that accumulated succinate during ischemia promotes the mitochondrial ROS burst during reperfusion at complex I by reverse electron transport (RET) (Chouchani et al., 2014). We therefore further measured RET-mediated H_2_O_2_ production in the presence of succinate in isolated cardiac mitochondria using the Amplex Red assay (Muller et al., 2004). Indeed, the rotenone-sensitive H_2_O_2_ production was significantly attenuated in TG mitochondria (Fig 5C), suggesting that NDUFAB1 is able to limit mitochondrial ROS production *via* the RET mechanism presumably through enhancing the assembly and activity of complex I and SCs.

**Figure 5.**
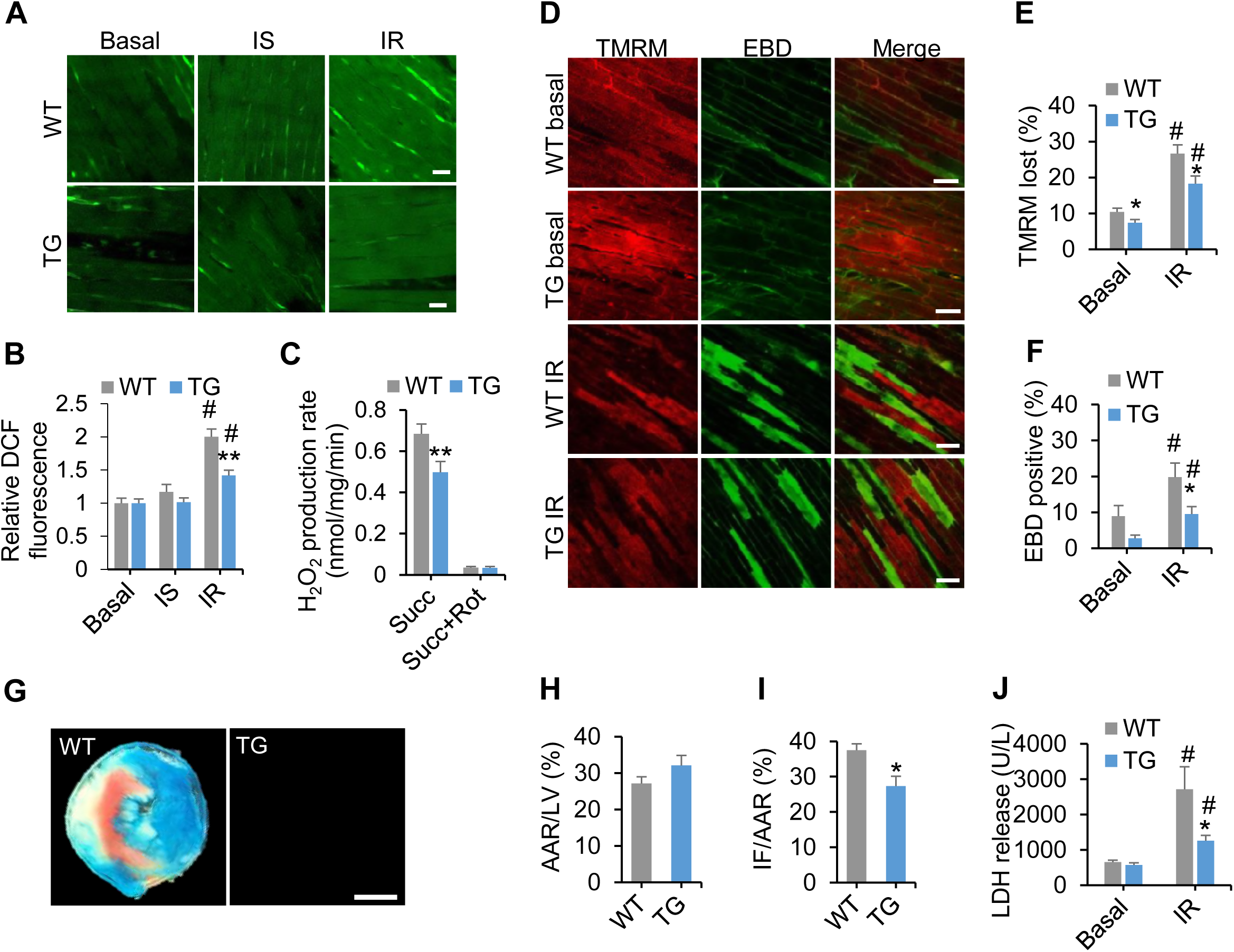
NDUFAB1 overexpression protects the heart against IR injury. TG and WT male littermates (2–4 months old) were used. **(A)** Alleviated myocardial ROS production in TG hearts. Confocal images of DCF-stained myocardium after Langendorff perfusion and IR injury. Ischemia (IS) was mimicked by stopping perfusion. See Figure EV7A for experimental protocol. Scale bars, 20 μm. **(B)** Statistics of **(A)** (mean ± s.e.m.; n = 32–51 image frames from 4–7 male mice per group; ^∗∗^***P*** <0.01 *versus* WT, ^#^***P*** <0.01 after IR *versus* basal). **(C)** Rotenone (Rot)-sensitive H_2_O_2_ production by RET at complex I in WT and TG cardiac mitochondria (mean ± s.e.m.; n = 6 male mice per group; ^∗∗^***P*** <0.01 *versus* WT). Succinate (Succ, 5 mM) was used as the substrate of complex II and H_2_O_2_ was measured with Amplex red. **(D)** Images for simultaneous measurement of ΔΨ_m_ (TMRM, red) and cardiomyocyte death (EBD uptake, green) in Langendorff-perfused hearts subjected to IR injury. Note that all cells with intact ΔΨ_m_ are EBD-negative, and all EBD-positive cells display loss of ΔΨ_m_, but not *vice versa.* Scale bars, 50 μm. **(E, F)** Percentages of cells with loss of ΔΨ_m_ **(E)** and EBD-positive cells **(F)** (mean ± s.e.m.; n = 9–10 male mice per group; ^∗^***P*** <0.05 *versus* WT, ^#^*P* <0.01 after IR *versus* basal). **(G)** Representative cross-sections of WT and TG hearts after IR injury. See Figure EV7B for the experimental protocol. White, infarcted area (IF); red, rest of the area at risk (AAR); blue, tissue not at risk. Scale bar, 1 mm. **(H, I)** Statistics of AAR **(H)** and IF **(I)** reported as the ratios of AAR to left ventricular area (LV) and IF to AAR (n = 5–8 male mice per group; ^∗^***P*** <0.05 *versus* WT). (J) Measurement of serum LDH release after IR (n = 10–13 male mice per group; ^∗^***P*** <0.05 *versus* WT, ^#^***P*** <0.01 after IR *versus* basal).

Because loss of ΔΨ_m_ not only manifests a defective bioenergetics but also is an early event of cell death (Matsumoto-Ida et al., 2006, Wang et al., 2010), we further assessed cardiac damage by monitoring loss of ΔΨ_m_ with TMRM and membrane integrity with Evans Blue dye (EBD). Our results showed that cells that both lost TMRM and were EBD-positive were significantly fewer in TG than WT hearts (Fig 5D-5F). Moreover, we adopted an *in vivo* IR experimental protocol in which a 30-min ischemia was followed by 24-h reperfusion (Fig EV7B). In agreement with the *ex vivo* IR results, the area of infarction was significantly smaller in TG than WT hearts (Fig 5G-5I). Meanwhile, LDH release induced by IR injury was significantly attenuated in TG mice (Fig 5J). Altogether, these findings demonstrate that increasing the abundance of mitochondrial NDUFAB1 is sufficient to protect the heart against IR injury.

## Discussion

The heart is quite sensitive to mitochondrial dysfunction such that most individuals with mutated mitochondrial proteins eventually develop cardiomyopathy (DiMauro & Schon, 2003). Not only are mitochondria the predominant powerhouses of the heart, supplying more than 90% of the ATP (Ingwall, 2002), but also excessive mitochondrial ROS are the culprits in many oxidative heart diseases including IR injury. Here we have shown, for the first time, that NDUFAB1 is a key endogenous regulator of mitochondrial bioenergetics in the mammalian heart and a cardio-protector when the heart is subjected to stress and injury. Our *ex vivo* and *in vivo* experimental results have established the essential role of NDUFAB1 in the maintenance of normal cardiac functions and the cardio-protective effects of NDUFAB1 overexpression. In the cKO heart, NDUFAB1 ablation led to a lowered ΔΨ_m_ and diminished ATP production. To aggravate the situation of an insufficient energy supply, more electrons from the ETC are diverted to the production of superoxide, elevating ROS to pathologically high levels. Cardiac performance deteriorates precipitously once the energy reserve is exhausted and the ATP level can no longer be held constant. De-compensatory myocardial remodeling also ensues, manifested as myocardial fibrosis, cardiomyocyte hypertrophy, and dilation of the ventricles with thinning of the ventricular walls, rapidly leading to heart failure and sudden death. Conversely and more importantly, increasing NDUFAB1 abundance makes the mitochondria not only more robust, featuring an augmented energy reserve capacity and efficiency, but also safer in terms of repressed ROS emission.

Both the improved bioenergetics and the diminished ROS production appear to be indispensable to NDUFAB1-mediated cardio-protection. On one hand, as the heart is highly demanding of energy, increasing the energy reserve capacity would make the heart more resistant to insults that tend to curtail ATP production. On the other hand, since excessive ROS constitute a crucial driving factor for IR injury in the heart (Eltzschig & Eckle, 2011, Murphy & Steenbergen, 2008, Yellon & Hausenloy, 2007), repression of basal ROS, and particularly the ROS burst during IR, should be beneficial. Specifically, we have found that NDUFAB1 overexpression represses RET-mediated ROS production at the onset of reperfusion after ischemia. Altogether, our results substantiate the emerging concept that energetically more robust and ROS-repressed mitochondria can protect the heart in stressful situations.

Another major finding in this study is that NDUFAB1 is a central coordinator in orchestrating the assembly of ETC complexes and SCs, mainly by regulating the biogenesis of FeS clusters (Cory et al., 2017, Van Vranken et al., 2016). In the absence of NDUFAB1, the assembly of complexes I-III, but not complex IV, was severely impaired. This result is in good agreement with the fact that complexes I-III all contain FeS clusters, but complex IV does not (Nicholls & Ferguson, 2002). Furthermore, only the FeS-containing subunits of complexes II and III were selectively down-regulated, and the premature complex III lacking the FeS-containing subunit UQCRFS1 accumulated in cKO mitochondria. For complex I, however, an additional mechanism must be invoked because all subunits examined, including those lacking FeS clusters, were downregulated in the absence of NDUFAB1. When NDUFAB1 was upregulated, greater effects were found on the assembly and enzymatic activity of complex I than complexes II or III. These differential effects are not unexpected, because cryo-electron microscopy has identified two copies of NDUFAB1 in complex I, and these are critically positioned for inter-subunit interactions within the complex (Fiedorczuk et al., 2016). Therefore, NDUFAB1 upregulation enhances complex I activity and assembly presumably through stabilizing inter-subunit interactions.

In addition to regulating ETC complex assembly, NDUFAB1 also plays an important role in the formation of SCs: its ablation halved while its overexpression significantly augmented the SC content. It is possible that NDUFAB1 modulates SC formation indirectly by regulating the assembly of individual complexes I-III, because it has been suggested that SCs form after complete assembly of the individual complexes (Guerrero-Castillo et al., 2017, Letts et al., 2016, Stroud et al., 2016). Nevertheless, our data cannot exclude the alternative possibility that NDUFAB1 participates directly in the assembly of SCs. If this is the case, NDUFAB1 might reversely regulate the abundance of individual complexes by affecting SC formation, since SCs have been suggested to stabilize individual complexes (Acin-Perez et al., 2004, Diaz et al., 2006, Milenkovic et al., 2017, Schagger et al., 2004). In agreement with the importance of SC formation in elevating mitochondrial electron transfer efficiency (Lopez-Fabuel et al., 2016, Maranzana et al., 2013, Milenkovic et al., 2017, Porras & Bai, 2015), the enhanced SCs in TG hearts constitute an important mechanism underlying NDUFAB1-mediated cardioprotection against injuries.

In summary, we have demonstrated that NDUFAB1 plays an essential role in mitochondrial bioenergetics and cardiac function through coordinating the assembly of individual ETC complexes and SCs. Augmenting NDUFAB1 abundance is sufficient to enhance the energy reserve capacity and efficiency of mitochondria, lower ROS production, and protect the heart against injurious insults. It is thus tempting to speculate that targeting NDUFAB1 to promote ETC complex and SC assembly might afford a potential therapeutic strategy for the treatment of mitochondrial bioenergetics-centered heart diseases.

## Materials and Methods

### Study approval

All animal experiments were carried out following the rules of the American Association for the Accreditation of Laboratory Animal Care International and the Guide for the Care and Use of Laboratory Animals published by the US National Institutes of Health (NIH Publication No. 85-23, revised 1996). All procedures were approved by the Animal Care Committee of Peking University accredited by AAALAC International (IMM-ChengHP-14).

### Generation of *Ndufab1* knockout and transgenic mice

Floxed *Ndufab1* mice were generated by standard techniques using a targeting vector containing a neomycin-resistance cassette flanked by Frt sites. Briefly, exon 3 of the *Ndufab1* gene was inserted into two flanking LoxP sites. After electroporation of the targeting vector into embryonic stem (ES) cells v6.5 (129×C57), G418-resistant ES cells were screened for homologous recombination by Southern blot. Two heterozygous recombinant ES clones were identified and microinjected into blastocysts from C57BL/6J mice to generate germline-transmitted floxed heterozygous mice (*Ndufab1*^f/+^). Homozygous Ndufab1-floxed mice (*Ndufab1*^f/f^) were obtained by inbreeding the Ndufab1^f/+^ mice.

Cardiac-specific *Ndufab1* knockout mice were generated as previously described (Zheng et al., 2009). Briefly, *Ndufab1*^f/f^ mice were bred with Mlc2v-Cre mice in which Cre recombinase expression was controlled by the myosin light chain 2v promoter to generate double heterozygous Mlc2v-Cre and *Ndufab1* floxed mice (Mlc2v-Crel Ndufab1^f/+^). The mice were then backcrossed with homozygous *Ndufab1*^f/f^ mice to generate Mlc2v-Cre^+^/*Ndufab1*^f/f^ cardiac-specific *Ndufab1* knockout mice and Mlc2v-Cre^-^/Ndufab1^f/f^ littermate controls. Mice were genotyped by PCR analysis using tail DNA, and flox primers (forward, ACAAATTCTCCCTGATGTCCTT; reverse, TGTTCAACTTCATTTTGAGGTGGT) and Cre primers (forward, GCGGTCTGGCAGTAAAAACTATC; reverse, GTGAAACAGCATTGCTGTCACTT) were used.

To generate global *Ndufab1* knockout mice, Ndufab1^f/f^ mice were bred with Prm-Cre mice in which Cre recombinase expression was controlled by the mouse protamine 1 promoter in the male germ line to generate heterozygous Prm-Cre and *Ndufab1* floxed mice (Prm-Cre^+^/Ndufab1^f/+^). The male mice were bred with wild-type (WT) C57BL/6J females to generate heterozygous *Ndufab1*^+/-^ mice. The heterozygous mice were intercrossed to generate homozygous *Ndufab1*^-/-^ mice. Mice were genotyped by PCR analysis using tail DNA, and WT primers (forward, ACAAATTCTCCCTGATGTCCTT; reverse, ACTGTATTCATGCCAGCATGTG) and KO primers (forward, ACAAATTCTCCCTGATGTCCTT; reverse, TCTGTTGTGCCCAGTCATAG) were used.

To generate pan-tissue transgenic mice expressing *Ndufab1*, the *Ndufab1* cDNA was cloned into the pUCCAGGS vector downstream of the chicken β-promoter. The construct was linearized with Hind III and Pvu I to release the transgenic cassette, purified with a DNA purification kit (Qiagen), and microinjected into fertilized eggs of C57BL/6J mice. The mice were genotyped by PCR analysis using the primers AGCCTCTGCTAACCATGTTC (forward) and GTCCAAACTGTCTAAGCCCA (reverse).

The mice were housed under a 12-h light cycle; food and water were provided *ad libitum.*

### Echocardiography

Mice were anesthetized with isoflurane (2% in 100% O2 at 0.5 L/min) and transthoracic echocardiography was performed using a VEVO-2100 Imaging System (Visual Sonics). The heart was imaged using M-mode and two-dimensional measurements were made at the level of the papillary muscles. Data represent the average of at least five separate scans in a random-blind fashion. Left ventricular posterior wall thickness at the end-diastolic and end-systolic phases (LVPWd and LVPWs), left ventricular internal diameter in end-diastole and end-systole (LVIDd and LVIDs), and end-diastolic and end-systolic interventricular septum thickness (IVSd and IVSs) were measured from the M-mode image. Fractional shortening (FS) and ejection fraction (EF) were used to assess systolic function.

### Masson trichrome staining

Mouse hearts were excised and fixed in 4% paraformaldehyde overnight, embedded in paraffin, and sectioned at 4 μm. Fibrosis was visualized by Masson trichrome staining. Ten fields were chosen randomly and imaged under an Olympus BX51 light microscope at ×20 magnification. ImageJ was used for fibrosis quantification.

### Transmission electron microscopy

Freshly-excised hearts were perfused with 1% glutaraldehyde and 4% paraformaldehyde for 30 min. After dissecting into 1-2 mm^3^ blocks, samples were immediately fixed with 2.5% glutaraldehyde and 4% paraformaldehyde, then post-fixed with 1% osmium tetroxide. After dehydration in a graded series of acetone, the samples were embedded in Spurr resin and sectioned with a glass knife on a Leica Ultracut R cutter. Ultra-thin sections (70–90 nm) were imaged with a transmission electron microscope (Tecnai G2 20 Twin, FEI). Mitochondrial area was calculated by analyzing images of mitochondria with a clear outer membrane on the micrographs using ImageJ freehand selection, and the number of pixels was converted to the actual mitochondrial cross-sectional area according to the magnification and pixel size of each micrograph. Mitochondrial volume fraction was analyzed by point-counting method (Medeiros, 2008). Briefly, a square lattice grid of 400 intersections was placed over each micrograph and the number of times a mitochondrion located at an intersection was counted. The final volume fraction was calculated by dividing the number with the total intersection number.

### Isolation of adult mouse cardiomyocytes

Single ventricular myocytes were enzymatically isolated from the hearts of WT, *Ndufab1* cardiac-specific knockout, or pan-tissue transgenic mice, as described previously (Cheng et al., 1993, Shang et al., 2016). Freshly-isolated cardiomyocytes were plated on laminin-coated (Sigma) culture dishes for 1 h and then the attached cells were maintained in Dulbecco’s modified Eagle’s medium (Invitrogen, Carlsbad, CA, USA) along with 10% FBS (Hyclone), 5 mM BDM (Sigma), and 1% insulin-transferrin-selenium supplement (Invitrogen) until use.

### Isolation and respiration measurement of cardiac mitochondria

Mouse hearts were washed with ice-cold isolation buffer (210 mM mannitol, 70 mM sucrose, 5 mM HEPES (pH 7.4), 1 mM EGTA, and 1 mg/ml BSA), minced, and homogenized. After the homogenate was centrifuged at 4°C for 10 min at 700 g, the supernatant was collected and further centrifuged at 4°C for 10 min at 12000 g. The pellet was re-suspended for functional assessment. The protein concentration of the mitochondrial preparation was determined with a NanoDrop Microvolume Spectrophotometer. 100 μg mitochondria was used for the respiration measurement.

Mitochondrial respiration was evaluated by measuring oxygen consumption with a Clark-type oxygen electrode (Strathkelvin 782 2-Channel Oxygen System version 1.0; Strathkelvin Instruments, Motherwell, UK). Briefly, isolated mitochondria were re-suspended in respiration buffer (in mM: 225 mannitol, 75 sucrose, 10 KCl, 10 Tris-HCl, 5 KH_2_PO_4_, pH 7.2) at 25°C with different substrates. State II and III respiration rates were each measured in the absence and presence of 100 μM ADP. The respiratory control ratio was defined as the ratio of the state III to the state II respiratory rate.

### BN-PAGE analysis of mitochondrial SCs and in-gel activity

BN-PAGE was conducted using the NativePAGE™ system (Invitrogen). Briefly, isolated cardiac mitochondria were solubilized by digitonin (4 g/g protein) for 15 min on ice, then centrifuged at 15,000 rpm at 4°C for 30 min. After centrifugation, the supernatants were collected and the protein concentration was determined by BCA analysis (ThermoFisher). Coomassie blue G-250 (Invitrogen) was added to the supernatant to obtain a dye/detergent mass ratio of 4/1 and then the protein was loaded into a 4-16% non-denaturing polyacrylamide gel (Invitrogen). After electrophoresis, proteins were transferred to a PVDF membrane and then probed with specific antibodies against subunits of complex I (NDUFB8), complex II (SDHA), complex III (UQCRFS1 and UQCRC1), complex IV (COX IV), and complex V (ATPB). Blots were visualized using secondary antibodies conjugated with IRDye (LI-COR, Lincoln, NE, USA) and an Odyssey imaging system (LI-COR). The anti-NDUFB8, anti-UQCRC1, or anti-COXIV immunoblot bands with high molecular weight were used to indicate SCs containing complex I, complex III, or complex IV.

For in-gel activity analysis, 2.5 mg/mL nitro blue tetrazolium and 0.5 mg/mL NADH in 2 mM Tris-HCl (pH 7.4) were added to the 4-16% non-denaturing polyacrylamide gel after electrophoresis and incubated for 15 min at 37°C. The reaction was stopped with 10% acetic acid. The activity was determined by analyzing the color development using the Odyssey imaging system (LI-COR).

### Whole-cell respiration analysis

Mitochondrial respiration was measured using a Seahorse XF24 Extracellular Flux Analyzer (Seahorse Bioscience, North Billerica, MA, USA) according to the manufacturer’s instructions. Briefly, isolated cardiomyocytes were seeded onto an XF24 microplate at 300 cells/well. The cellular oxygen consumption rate (OCR) was monitored in unbuffered assay medium (Sigma D5030) supplemented with (in mM) 2 GlutaMAX (Gibco), 2.5 sodium pyruvate, and 25 glucose (pH 7.4 at 37°C), following the sequential addition of oligomycin (1 μM), carbonyl cyanide 4-(trifluoromethoxy) phenylhydrazone (FCCP, 500 nM), and rotenone (1 μM) and antimycin A (AA, 1 μM). Basal OCR refers to the respiration rate measured prior to the addition of oligomycin. ATP-coupled OCR was calculated by subtracting the OCR in the presence of oligomycin from basal OCR. Maximal OCR was calculated by subtracting the OCR in the presence of rotenone and AA from that in the presence of FCCP. Proton leak-coupled OCR was calculated by subtracting the OCR in the presence of rotenone and AA from that in the presence of oligomycin.

### Measurement of H_2_O_2_ production

H_2_O_2_ production in isolated mitochondria was measured using the Amplex^®^ Red hydrogen peroxide kit (Molecular Probes). Briefly, isolated mitochondria were incubated in an assay medium with 10 mM succinate, 50 mM Amplex red, and 5 units/ml HRP in each well of a 96-well plate. The increase in Amplex red fluorescence was followed over 60 min at room temperature with excitation at 530 nm and emission at >590 nm in a 96-well microplate reader (Biotek, Winooski, VT, USA). H_2_O_2_ production was indexed by the increase of Amplex red fluorescence per min.

### ATP measurement

The heart was excised and lysed with Tris-phenol. The myocardial ATP content was measured by luciferase assay according to manufacturer’s instructions (Promega, Madison, WI, USA).

### *In vivo* and *ex vivo* myocardial IR models

For *in vivo* IR surgery, 2–4-month-old male mice were anesthetized with avertin (2.5%, 0.01 mL/g, i.p.) and ventilated *via* a tracheostomy on a Minivent respirator. A midline sternotomy was performed, and a reversible coronary artery snare occluder was placed around the left anterior descending coronary artery. Myocardial IR was induced by tightening the snare for 30 min and then loosening it for 24 h. To measure the infarct size, the animals were anesthetized with avertin (2.5%, 0.015 mL/g, i.p.) and heparinized (400 USP U/kg, i.p.) at the end-point. The heart was excised and the ascending aorta was cannulated (distal to the sinus of Valsalva), then perfused retrogradely with Alcian blue dye (1%) to visualize the area not at risk. The coronary artery was re-occluded at the site of occlusion before perfusion with Alcian blue. Then, the heart was frozen at -80 °C for 10 min and cut into 1 mm slices (5-6 slices per heart), which were then incubated in sodium phosphate buffer containing 1.5% 2,3,5-triphenyl-tetrazolium chloride for 15 min to visualize the unstained infarcted region. Infarct area, area at risk (AAR), and left ventricular area were determined by planimetry using ImageJ. The infarct size was calculated as infarct area divided by AAR. For lactate dehydrogenase (LDH) measurement, blood samples were collected before surgery and after 24 h of reperfusion, and centrifuged for 10 min at 3,000 r.p.m. to obtain serum. LDH was spectrophotometrically assayed using a kit from Jingyuan Biology (Shanghai, China).

For *ex vivo* IR, the heart was excised from male mice (2-4 months old), and the ascending aorta was cannulated with a customized needle. The heart was perfused in the Langendorff configuration under constant perfusion pressure (1000 mm H_2_O) with oxygenated (100% O_2_) Tyrode’s solution (pH 7.4 at 37°C). The Langendorff-perfused heart was placed on the stage of the confocal microscope and images were captured ~30 μm deep into the epimyocardium of the left ventricle. Motion artifacts due to spontaneous beating of the heart (~480/min) were minimized using 10 μM blebbistatin (Sigma). After a 10-min stabilization period, the heart was perfused with 15 μM DCF (an indicator of ROS) or 500 nM TMRM (an indicator of mitochondrial membrane potential) and 0.05% EBD (an indicator of plasma membrane integrity) for 20 min. The IR protocol used was as described previously (Matsumoto-Ida et al., 2006, Wang et al., 2010). Ischemia (30 min) was achieved by clamping the perfusion line and reperfusion (30 min) by releasing the clamp. Confocal images were captured from multiple, randomly selected regions prior to, during ischemia, and after the IR, at excitation wavelengths of 488, 543, and 633 nm and emission wavelengths of 500-740, 565-595, and >650 nm for DCF, TMRM, and EBD, respectively.

### Confocal microscopy

An inverted confocal microscope (Zeiss LSM 710) with a 40×, 1.3 NA oil-immersion objective was used for imaging. For mitoSOX measurement, the indicator (5 μM) was loaded into isolated cardiomyocytes at 37°C for 30 min followed by 3 washes with Tyrode’s solution. The mitoSOX fluorescence was measured by excitation at 514 nm and emission at 580–740 nm. To measure mitochondrial membrane potential in isolated cardiomyocytes, TMRM (50 nM) was loaded at 37°C for 10 min followed by 3 washes and the fluorescence was measured by excitation at 543 nm and emission at >560 nm.

### RNA-seq analysis

Using TRIzol reagent (Invitrogen), total RNA was isolated from left ventricular myocardium in cKO and WT male mice (2-4 months old) according to the manufacture’s protocol. The quality of the extracted RNA was controlled using Agilent 2100. A sequencing library was prepared using the NEBNext^®^Ultra^TM^ RNA Library Prep Kit for Illumina^®^ (New England Biolabs, Ipswich, MA, USA) following the manufacturer’s recommendations. Deep sequencing was then performed on an Illumina Hiseq 4000 platform and 150 bp paired-end reads were generated. To obtain the differentially-expressed genes, RNA-seq reads were first aligned to the mouse genome (genome version mm9) with tophat2 (version 2.1.1; main parameters: ‐‐read-mismatches 8, ‐‐min-anchor 8, ‐‐segment-length 30, ‐‐segment-mismatches 2) and all the mapping results were evaluated as previously described (Zhang et al., 2017). DEseq2 packages were used to obtain differentially-expressed genes with a cutoff of fold-change in parameP-value <0.05. The functional annotation and pathway enrichment analysis for both upregulated and downregulated genes were performed using Database for Annotation, Visualization, and Integrated Discovery(Huang da et al., 2009a, Huang da et al., 2009b) with a P-value cutoff of 0.05 under the Benjamini test.

### Western-blot analysis

Frozen myocardial tissue or isolated mitochondria was homogenized with denaturing lysis buffer, and 10-50 μg of protein per sample was separated on 12% SDS-PAGE. After electrophoresis, proteins were transferred to a PVDF membrane and then probed with specific antibodies. Blots were visualized using secondary antibodies conjugated with IRDye (LI-COR) and an Odyssey imaging system (LI-COR). The antibodies for NDUFAB1 (TA308227), SDHB (TA351639), and SDHD (TA326622) were from Origene, those for NDUFB8 (ab110242), NDUFS4 (ab137064), SDHA (ab14715), UQCRC1 (ab110252), UQCRB (ab190360), ATPB (ab14730), and ATP5A (ab176569) from Abcam, and those for NDUFS1 (12444-1-AP), NDUFS6 (14417-1-AP), NDUFS7 (15728-1-AP), UQCRFS1 (18443-1-AP), and COXIV (11242-1-AP) from ProteinTech.

### RNA isolation and real-time PCR (RT-PCR)

Total RNA was isolated from mouse ventricular myocardium with TRIzol reagent (Invitrogen) according to the manufacturer’s instructions, and then converted to cDNA using TransScript One-Step gDNA Removal and cDNA Synthesis Mix (Transgen Biotech). Quantitative RT-PCR reactions were performed using Trans Start Green qPCR Super Mix (TransGen Biotech) and a BioRad CFX96 Touch RT-PCR Detection System. The primers used were as follows:

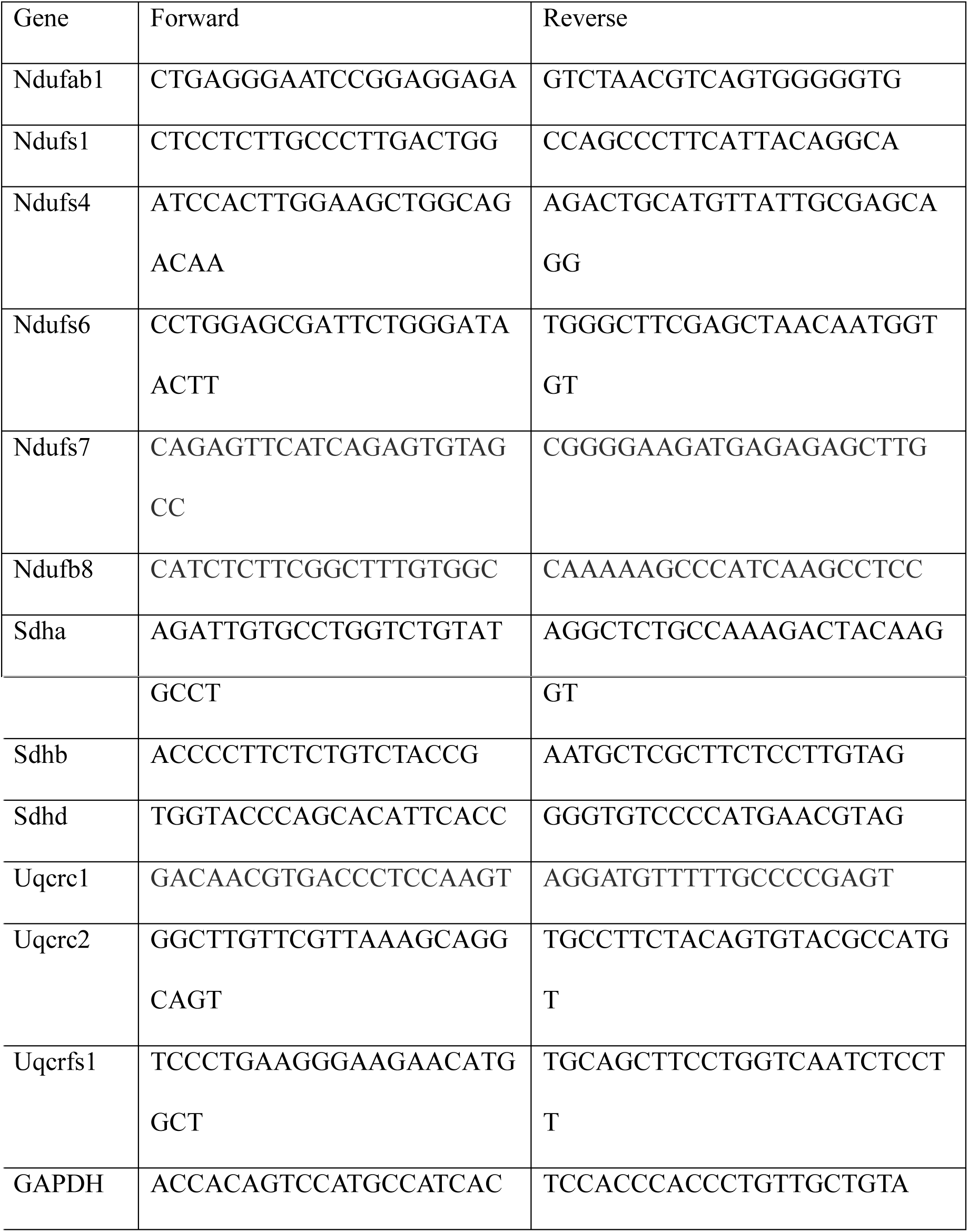

### Statistics

Data are expressed as mean ± SEM. When appropriate, Student’s t-test was applied to determine statistical significance. The log-rank test was used for survival curves. P <0.05 was considered statistically significant.

## Footnotes

**Author contributions** TH, HC, and XW conceived and designed the study. TH, CJ, and XH developed the methodology. TH, RZ, YW and QM conducted experiments. TH, HC, and XW wrote the manuscript.

## Acknowledgements

We thank Ms. Yuli Liu for echocardiography, Dr. Lin Pan and Ms. Dongwei Ma for histology, Drs. Liping Wei and Chuanyun Li for bioinformatics analysis, Dr. Rongli Zhang for IR experiments, and Drs. Iain C. Bruce, Ruiping Xiao, Yingxian Li, and Yan Zhang for valuable comments.

## Sources of founding

This work was supported by the National Key Basic Research Program of China (2017YFA0504000, 2016YFA0500403, and 2013CB531200) and the National Science Foundation of China (31470811, 31670039, and 31521062).

## Figure legend

**Figure EV1.**
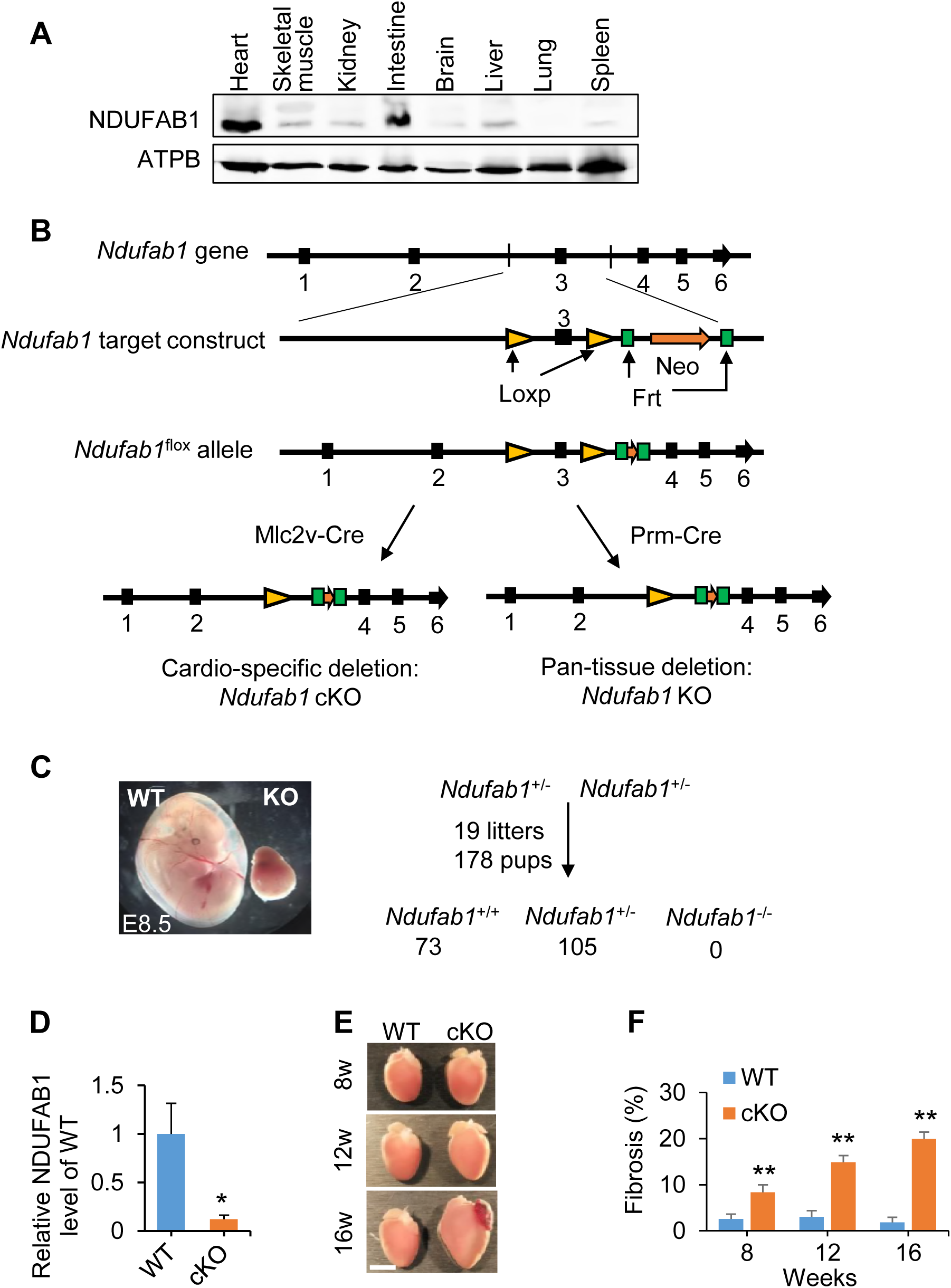
Generation and phenotyping of *Ndufab1* cardiac-specific knockout (cKO) or pan-tissue knockout (KO) mouse models. **(A)** Western blots for *Ndufab1* expression in different mouse organs. Anti-ATPB served as the loading control. **(B)** Schematic of gene-targeting strategy. LoxP sites flank exon 3 of the *Ndufab1* gene and Frt sites flank the neomycin resistance (Neo) cassette. *Ndufab1*^flox/flox^ mice were crossed with Mlc2v-Cre or Prm-Cre mice to allow cardiomyocyte-specific or pan-tissue deletion of *Ndufab1*, respectively. **(C)** Embryonic lethality of pan-tissue knockout of *Ndufab1.* Left, representative images of E8.5 wild-type (WT) and KO embryos. Right, summary of live pups recovered from Ndufab1+^/-^ intercrossing. **(D)** Relative expression of NDUFAB1 in WT (flox/flox, Mlc2v-Cre^-^) and cKO (flox/flox, Mlc2v-Cre^+^) hearts (mean ± s.e.m.; n = 4 mice per group; ^∗^***P*** <0.05 *versus* WT). **(E)** Representative images of gross heart morphology at 8-, 12-, and 16-weeks (w) of age. Note that a blood clot has pooled in the left atrial appendage of the 16-week old cKO heart (scale bar, 5 mm). **(F)** Statistics of fibrosis (as shown in Figure 1G) (mean ± s.e.m.; n = 24–45 fields from 3 hearts per group; ^∗∗^ ***P*** <0.01 *versus* WT).

**Figure EV2.**
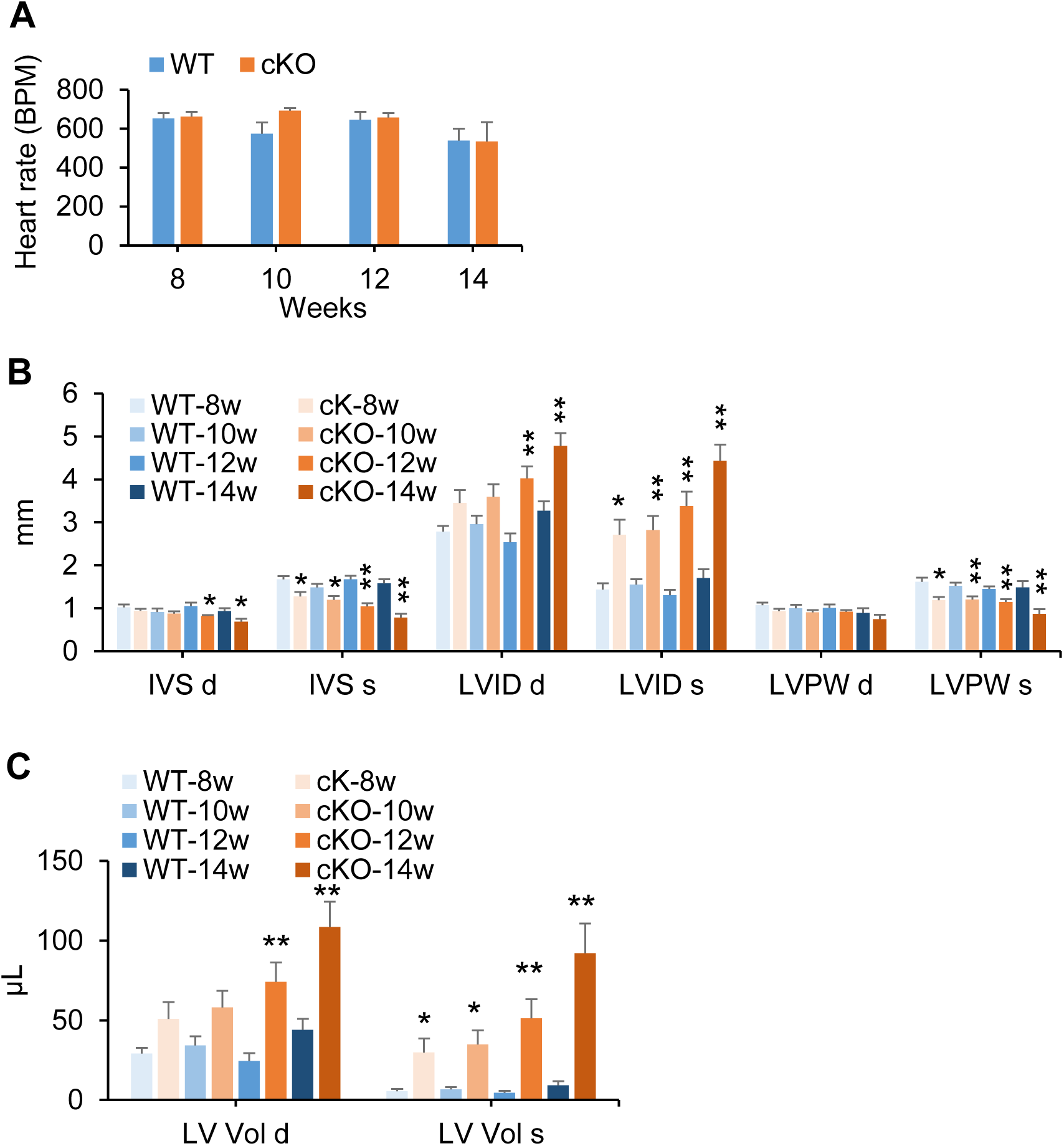
Echocardiographic analysis of cardiac functions of WT and cKO mice at different ages. **(A)** Heart rate (BPM, beats per minute; mean ± s.e.m.; n = 3–10 mice per group). **(B)** Interventricular septum thickness at end-diastole and end-systole (IVS d, s), left ventricular internal diameter at end-diastole and end-systole (LVID d, s), and left ventricular posterior wall thickness at end-diastole and end-systole (LVPW d, s) (mean ± s.e.m.; n = 3–9 mice per group; ^∗^ ***P*** <0.05, ^∗∗^ ***P*** <0.01 *versus* WT). **(C)** Left ventricular volume at end-diastole and end-systole (LV Vol d, s) (mean ± s.e.m.; n = 3–9 mice per group; ^∗^ ***P*** <0.05, ^∗∗^ ***P*** <0.01 *versus* WT).

**Figure EV3.**
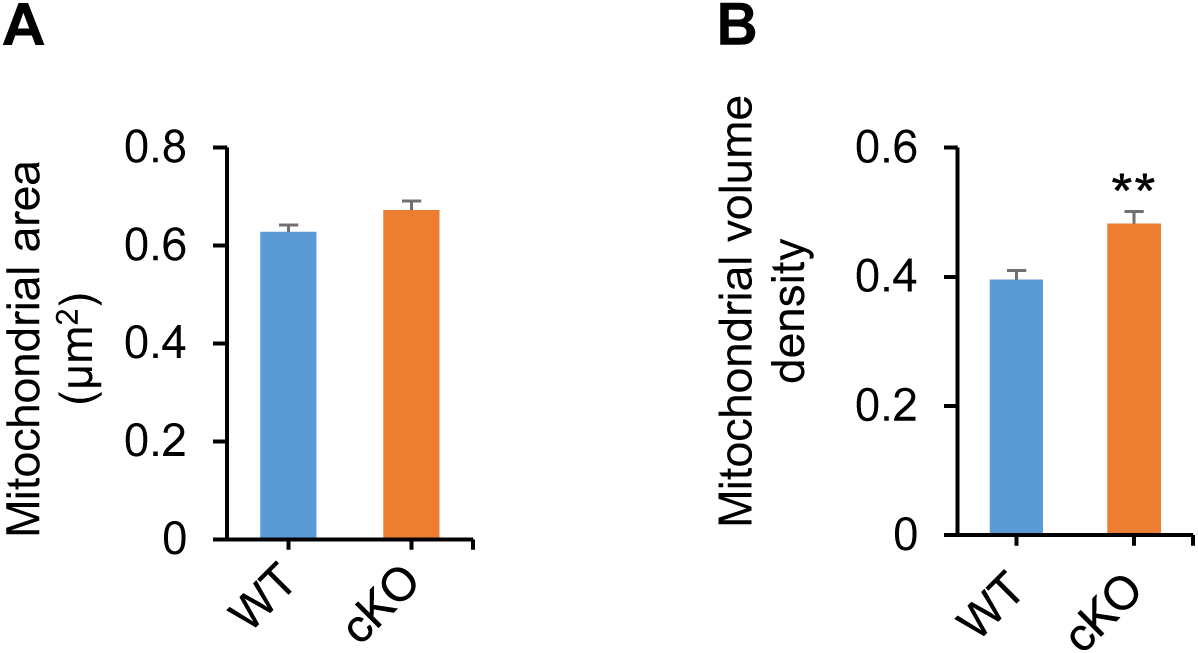
Myocardial mitochondrial area and volume density quantified from electron micrographs (as shown in Figure 2A) (mean ± s.e.m.; n = 157–172 mitochondria for **(A)** and 18–42 images for **(B)**; 3 mice per group; ^∗∗^ ***P*** <0.01 *versus* WT).

**Figure EV4.**
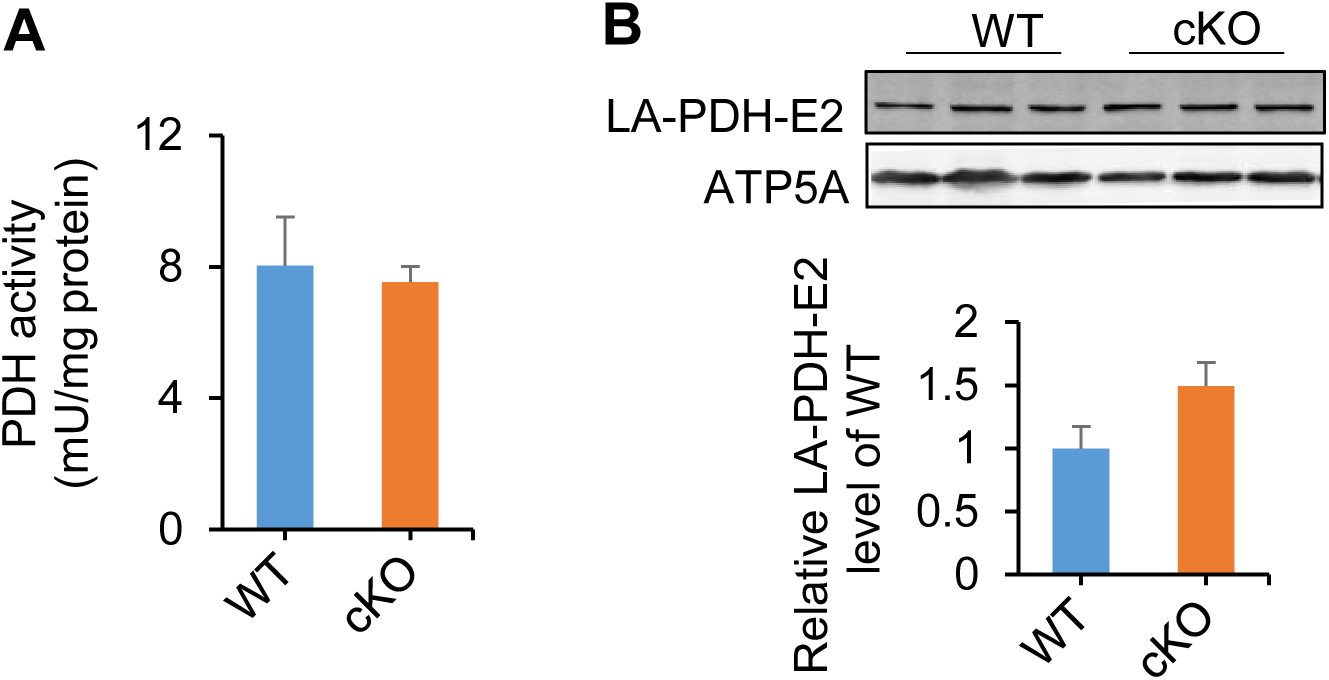
Pyruvate dehydrogenase (PDH) activity (A) and lipoic acid (LA)-conjugated E2 of PDH complex expression (B) in WT and cKO hearts. Data are mean ± s.e.m.. n = 3 mice per group.

**Figure EV5.**
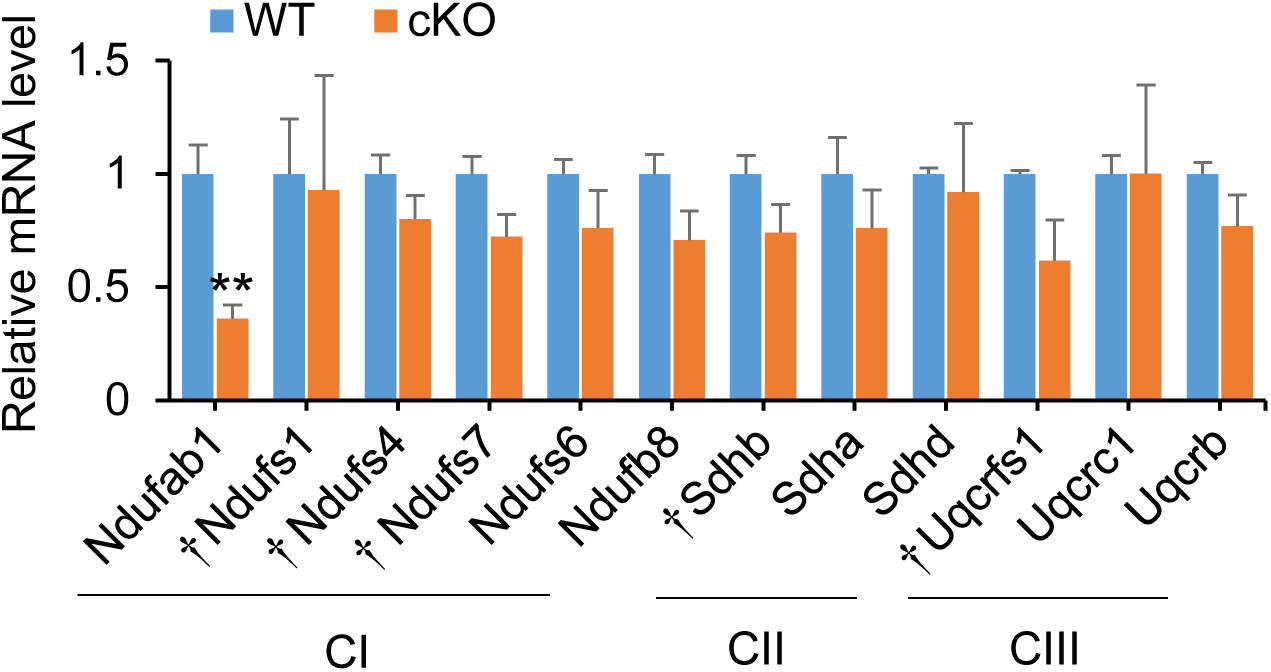
Real-time PCR analysis of subunit expression of complexes I-III (CI-CIII) (mean ± s.e.m.; n = 4–6 independent experiments per group; ^∗∗^ ***P*** <0.01 *versus* WT; ^†^FeS-containing proteins).

**Figure EV6.**
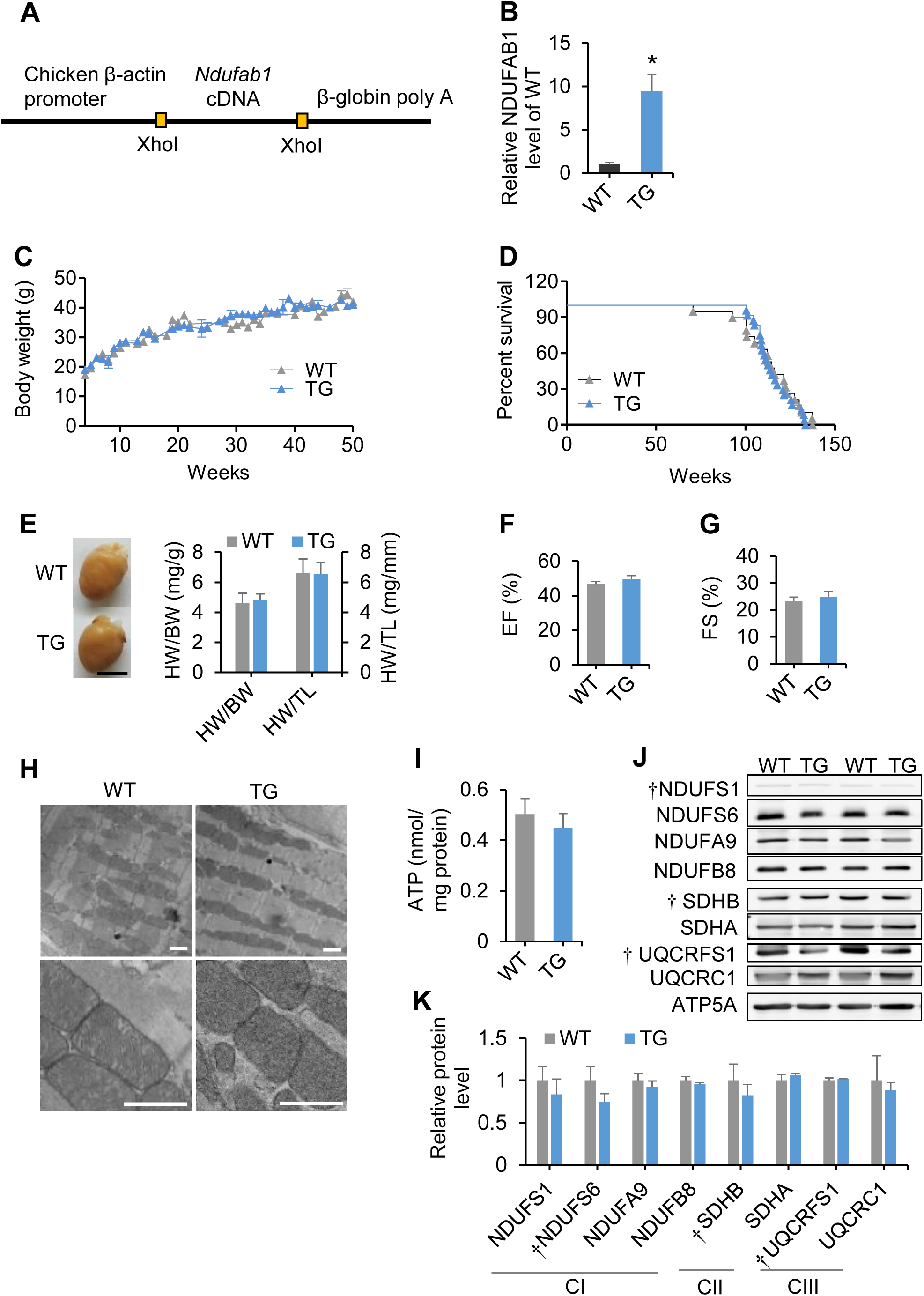
Generation and phenotyping of *Ndufab1* transgenic (TG) mice. **(A)** Schematic of transgenic construct for generating TG mice. Mouse *Ndufab1* cDNA was cloned into the pUCCAGGS vector downstream of the chicken β-actin promoter. **(B)** NDUFAB1 in WT and TG hearts (mean ± s.e.m.; n = 3 mice per group; ^∗^ ***P*** <0.05 *versus* WT). **(C)** Growth curves of WT and TG mice (n = 3–15 mice per time point). **(D)** Kaplan-Meier survival curves of WT and TG mice (n = 19 for WT and 24 for TG mice). **(E)** Cardiac morphology. Left panel, representative photographs of WT and TG hearts (scale bar, 5 mm). Right panel, ratios of heart weight (HW) to body weight (BW) or tibial length (TL) (mean ± s.e.m.; n = 7–36 mice per group). The mice were 12 weeks old. **(F, G)** Echocardiographic analysis of cardiac function (EF, ejection fraction; FS, fractional shortening; mean ± s.e.m.; n = 6 mice per group). The mice were 8–12 weeks old. **(H)** Electron micrographs of left ventricular tissue (scale bars, 1 μm). (I) ATP content of WT and TG hearts (mean ± s.e.m.; n = 5–6 mice per group). The mice were 8–12 weeks old. (J) Western blots for subunits of complexes I-III (^†^FeS-containing subunits). (K) Statistics of **(J)** (mean ± s.e.m.; n = 3 independent experiments per group).

**Figure EV7.**
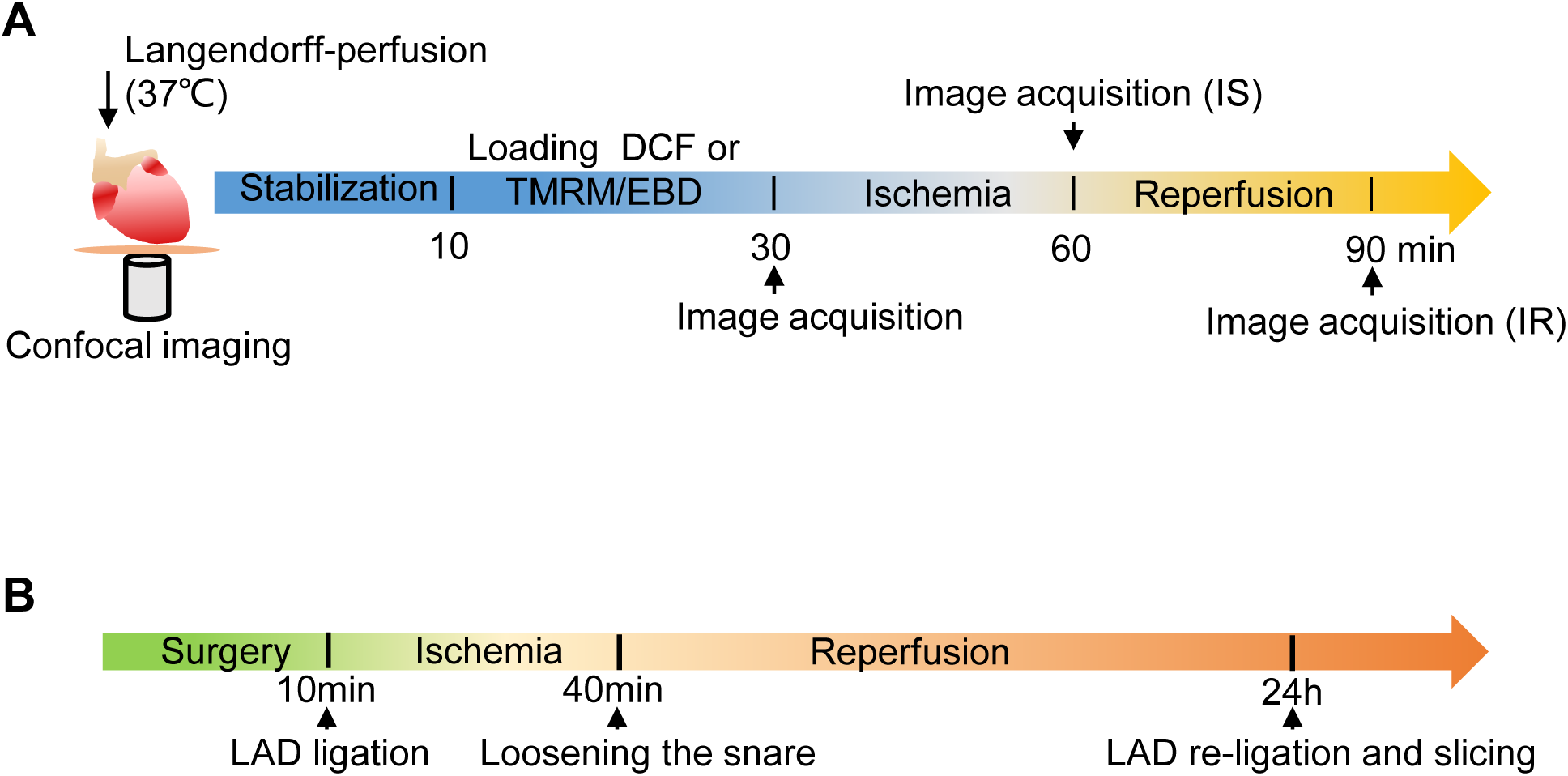
Schematic of *ex vivo* and *in vivo* IR protocols. **(A)** Schematic of the *ex vivo* IR protocol for the Langendorff-perfused heart. Briefly, the heart was subjected to ischemia for 30 min and then reperfusion for 30 min. Images were captured before ischemia, at 30 min of ischemia (IS), and at 30 min of reperfusion (IR). **(B)** Schematic of the *in vivo* IR protocol. A reversible coronary artery snare occluder was placed around the left anterior descending (LAD) coronary artery of the anesthetized mouse. Myocardial I/R was induced by tightening the snare for 30 min and then loosening it for 24 h.

## References

Acin-Perez R, Bayona-Bafaluy MP, Fernandez-Silva P, Moreno-Loshuertos R, Perez-Martos A, Bruno C, Moraes CT, Enriquez JA (2004) Respiratory complex III is required to maintain complex I in mammalian mitochondria. Molecular cell 13: 805–15

Acin-Perez R, Enriquez JA (2014) The function of the respiratory supercomplexes: the plasticity model. Biochimica et biophysica acta 1837: 444–50

Allue I, Gandelman O, Dementieva E, Ugarova N, Cobbold P (1996) Evidence for rapid consumption of millimolar concentrations of cytoplasmic ATP during rigor-contracture of metabolically compromised single cardiomyocytes. The Biochemical journal 319 (Pt 2): 463–9

Balaban RS, Kantor HL, Katz LA, Briggs RW (1986) Relation between work and phosphate metabolite in the in vivo paced mammalian heart. Science 232: 1121–3

Beckman KB, Ames BN (1998) The free radical theory of aging matures. Physiological reviews 78: 547–81

Cheng H, Lederer WJ, Cannell MB (1993) Calcium sparks: elementary events underlying excitation-contraction coupling in heart muscle. Science 262: 740–4

Chouchani ET, Pell VR, Gaude E, Aksentijevic D, Sundier SY, Robb EL, Logan A, Nadtochiy SM, Ord ENJ, Smith AC, Eyassu F, Shirley R, Hu CH, Dare AJ, James AM, Rogatti S, Hartley RC, Eaton S, Costa ASH, Brookes PS et al. (2014) Ischaemic accumulation of succinate controls reperfusion injury through mitochondrial ROS. Nature 515: 431–435

Cory SA, Van Vranken JG, Brignole EJ, Patra S, Winge DR, Drennan CL, Rutter J, Barondeau DP (2017) Structure of human Fe-S assembly subcomplex reveals unexpected cysteine desulfurase architecture and acyl-ACP-ISD11 interactions. Proceedings of the National Academy of Sciences of the United States of America 114: E5325–E5334

Diaz F, Fukui H, Garcia S, Moraes CT (2006) Cytochrome c oxidase is required for the assembly/stability of respiratory complex I in mouse fibroblasts. Molecular and cellular biology 26: 4872–81

DiMauro S, Schon EA (2003) Mitochondrial respiratory-chain diseases. The New England journal of medicine 348: 2656–68

Droge W (2002) Free radicals in the physiological control of cell function. Physiological reviews 82: 47–95

Eltzschig HK, Eckle T (2011) Ischemia and reperfusion‐‐from mechanism to translation. Nat Med 17: 1391–401

Feng D, Witkowski A, Smith S (2009) Down-regulation of mitochondrial acyl carrier protein in mammalian cells compromises protein lipoylation and respiratory complex I and results in cell death. The Journal of biological chemistry 284: 11436–45

Fiedorczuk K, Letts JA, Degliesposti G, Kaszuba K, Skehel M, Sazanov LA (2016) Atomic structure of the entire mammalian mitochondrial complex I. Nature 538: 406–410

Floyd BJ, Wilkerson EM, Veling MT, Minogue CE, Xia C, Beebe ET, Wrobel RL, Cho H, Kremer LS, Alston CL, Gromek KA, Dolan BK, Ulbrich A, Stefely JA, Bohl SL, Werner KM, Jochem A, Westphall MS, Rensvold JW, Taylor RW et al. (2016) Mitochondrial Protein Interaction Mapping Identifies Regulators of Respiratory Chain Function. Molecular cell 63: 621–632

Greggio C, Jha P, Kulkarni SS, Lagarrigue S, Broskey NT, Boutant M, Wang X, Conde Alonso S, Ofori E, Auwerx J, Canto C, Amati F (2017) Enhanced Respiratory Chain Supercomplex Formation in Response to Exercise in Human Skeletal Muscle. Cell metabolism 25: 301–311

Gu J, Wu M, Guo R, Yan K, Lei J, Gao N, Yang M (2016) The architecture of the mammalian respirasome. Nature 537: 639–43

Guaras A, Perales-Clemente E, Calvo E, Acin-Perez R, Loureiro-Lopez M, Pujol C, Martinez-Carrascoso I, Nunez E, Garcia-Marques F, Rodriguez-Hernandez MA, Cortes A, Diaz F, Perez-Martos A, Moraes CT, Fernandez-Silva P, Trifunovic A, Navas P, Vazquez J, Enriquez JA (2016) The CoQH2/CoQ Ratio Serves as a Sensor of Respiratory Chain Efficiency. Cell reports 15: 197–209

Guerrero-Castillo S, Baertling F, Kownatzki D, Wessels HJ, Arnold S, Brandt U, Nijtmans L (2017) The Assembly Pathway of Mitochondrial Respiratory Chain Complex I. Cell metabolism 25: 128–139

Guo R, Zong S, Wu M, Gu J, Yang M (2017) Architecture of Human Mitochondrial Respiratory Megacomplex I2III2IV2. Cell 170: 1247–1257

Hiltunen JK, Autio KJ, Schonauer MS, Kursu VA, Dieckmann CL, Kastaniotis AJ (2010a) Mitochondrial fatty acid synthesis and respiration. Biochimica et biophysica acta 1797: 1195–202

Hiltunen JK, Chen Z, Haapalainen AM, Wierenga RK, Kastaniotis AJ (2010b) Mitochondrial fatty acid synthesis‐‐an adopted set of enzymes making a pathway of major importance for the cellular metabolism. Prog Lipid Res 49: 27–45

Huang da W, Sherman BT, Lempicki RA (2009a) Bioinformatics enrichment tools: paths toward the comprehensive functional analysis of large gene lists. Nucleic acids research 37: 1–13

Huang da W, Sherman BT, Lempicki RA (2009b) Systematic and integrative analysis of large gene lists using DAVID bioinformatics resources. Nature protocols 4: 44–57

Ingwall JS (2002) ATP and the Heart. Norwell, Massachusetts: Kluwer Academic Publishers,

Ingwall JS (2004) Transgenesis and cardiac energetics: new insights into cardiac metabolism. Journal of molecular and cellular cardiology 37: 613–23

Jian C, Xu F, Hou T, Sun T, Jinghang L, Cheng H, Wang X (2017) Deficiency of PHB complex impairs respiratory supercomplex formation and activates mitochondrial flashes. J Cell Sci 130: 2620–2630

Lapuente-Brun E, Moreno-Loshuertos R, Acin-Perez R, Latorre-Pellicer A, Colas C, Balsa E, Perales-Clemente E, Quiros PM, Calvo E, Rodriguez-Hernandez MA, Navas P, Cruz R, Carracedo A, Lopez-Otin C, Perez-Martos A, Fernandez-Silva P, Fernandez-Vizarra E, Enriquez JA (2013) Supercomplex assembly determines electron flux in the mitochondrial electron transport chain. Science 340: 1567–70

Letts JA, Fiedorczuk K, Sazanov LA (2016) The architecture of respiratory supercomplexes. Nature 537: 644–648

Lopez-Fabuel I, Le Douce J, Logan A, James AM, Bonvento G, Murphy MP, Almeida A, Bolanos JP (2016) Complex I assembly into supercomplexes determines differential mitochondrial ROS production in neurons and astrocytes. Proceedings of the National Academy of Sciences of the United States of America 113: 13063–13068

Maack C, O’Rourke B (2007) Excitation-contraction coupling and mitochondrial energetics. Basic research in cardiology 102: 369–92

Maranzana E, Barbero G, Falasca AI, Lenaz G, Genova ML (2013) Mitochondrial respiratory supercomplex association limits production of reactive oxygen species from complex I. Antioxidants & redox signaling 19: 1469–80

Matsumoto-Ida M, Akao M, Takeda T, Kato M, Kita T (2006) Real-time 2-photon imaging of mitochondrial function in perfused rat hearts subjected to ischemia/reperfusion. Circulation 114: 1497–503

Matthews PM, Bland JL, Gadian DG, Radda GK (1981) The steady-state rate of ATP synthesis in the perfused rat heart measured by 31P NMR saturation transfer. Biochem Biophys Res Commun 103: 1052–9

Medeiros DM (2008) Assessing mitochondria biogenesis. Methods 46: 288–94

Milenkovic D, Blaza JN, Larsson NG, Hirst J (2017) The Enigma of the Respiratory Chain Supercomplex. Cell metabolism 25: 765–776

Mitsopoulos P, Chang YH, Wai T, Konig T, Dunn SD, Langer T, Madrenas J (2015) Stomatin-like protein 2 is required for in vivo mitochondrial respiratory chain supercomplex formation and optimal cell function. Molecular and cellular biology 35: 1838–47

Muller FL, Liu Y, Van Remmen H (2004) Complex III releases superoxide to both sides of the inner mitochondrial membrane. The Journal of biological chemistry 279: 49064–73

Murphy E, Steenbergen C (2008) Mechanisms underlying acute protection from cardiac ischemia-reperfusion injury. Physiol Rev 88: 581–609

Neely JR, Rovetto MJ, Whitmer JT, Morgan HE (1973) Effects of ischemia on function and metabolism of the isolated working rat heart. Am J Physiol 225: 651–8

Neubauer S (2007) The failing heart‐‐an engine out of fuel. The New England journal of medicine 356: 1140–51

Nicholls DG, Ferguson SL (2002) Bioenergetics. Academic Press,

Porras CA, Bai Y (2015) Respiratory supercomplexes: plasticity and implications. Front Biosci (Landmark Ed) 20: 621–34

Rosca MG, Vazquez EJ, Kerner J, Parland W, Chandler MP, Stanley W, Sabbah HN, Hoppel CL (2008) Cardiac mitochondria in heart failure: decrease in respirasomes and oxidative phosphorylation. Cardiovasc Res 80: 30–9

Runswick MJ, Fearnley IM, Skehel JM, Walker JE (1991) Presence of an acyl carrier protein in NADH:ubiquinone oxidoreductase from bovine heart mitochondria. FEBS letters 286: 121–4

Schagger H, de Coo R, Bauer MF, Hofmann S, Godinot C, Brandt U (2004) Significance of respirasomes for the assembly/stability of human respiratory chain complex I. The Journal of biological chemistry 279: 36349–53

Shang W, Gao H, Lu F, Ma Q, Fang H, Sun T, Xu J, Ding Y, Lin Y, Wang Y, Wang X, Cheng H, Zheng M (2016) Cyclophilin D regulates mitochondrial flashes and metabolism in cardiac myocytes. J Mol Cell Cardiol 91: 63–71

Sousa JS, Mills DJ, Vonck J, Kuhlbrandt W (2016) Functional asymmetry and electron flow in the bovine respirasome. eLife 5

Stroud DA, Surgenor EE, Formosa LE, Reljic B, Frazier AE, Dibley MG, Osellame LD, Stait T, Beilharz TH, Thorburn DR, Salim A, Ryan MT (2016) Accessory subunits are integral for assembly and function of human mitochondrial complex I. Nature 538: 123–126

Van Vranken JG, Jeong MY, Wei P, Chen YC, Gygi SP, Winge DR, Rutter J (2016) The mitochondrial acyl carrier protein (ACP) coordinates mitochondrial fatty acid synthesis with iron sulfur cluster biogenesis. Elife 5

Vinothkumar KR, Zhu J, Hirst J (2014) Architecture of mammalian respiratory complex I. Nature 515: 80–84

Wagener N, Ackermann M, Funes S, Neupert W (2011) A pathway of protein translocation in mitochondria mediated by the AAA-ATPase Bcs1. Molecular cell 44: 191–202

Wang X, Xie W, Zhang Y, Lin P, Han L, Han P, Wang Y, Chen Z, Ji G, Zheng M, Weisleder N, Xiao RP, Takeshima H, Ma J, Cheng H (2010) Cardioprotection of ischemia/reperfusion injury by cholesterol-dependent MG53-mediated membrane repair. Circulation research 107: 76–83

Wang X, Zhang X, Wu D, Huang Z, Hou T, Jian C, Yu P, Lu F, Zhang R, Sun T, Li J, Qi W, Wang Y, Gao F, Cheng H (2017) Mitochondrial flashes regulate ATP homeostasis in the heart. Elife 6

Wu M, Gu J, Guo R, Huang Y, Yang M (2016) Structure of Mammalian Respiratory Supercomplex I1III2IV1 Cell 167: 1598–1609

Yellon DM, Hausenloy DJ (2007) Myocardial reperfusion injury. The New England journal of medicine 357: 1121–35

Zhang SJ, Wang C, Yan S, Fu A, Luan X, Li Y, Sunny Shen Q, Zhong X, Chen JY, Wang X, Chin-Ming Tan B, He A, Li CY (2017) Isoform Evolution in Primates through Independent Combination of Alternative RNA Processing Events. Molecular biology and evolution 34: 2453–2468

Zheng M, Cheng H, Li X, Zhang J, Cui L, Ouyang K, Han L, Zhao T, Gu Y, Dalton ND, Bang ML, Peterson KL, Chen J (2009) Cardiac-specific ablation of Cypher leads to a severe form of dilated cardiomyopathy with premature death. Human molecular genetics 18: 701–13

